# The Antitumor Activities of Anti-CD47 Antibodies Require Fc-FcγR interactions

**DOI:** 10.1101/2023.06.29.547082

**Authors:** Juan C. Osorio, Patrick Smith, David A. Knorr, Jeffrey V. Ravetch

## Abstract

While anti-CD47 antibodies hold promise for cancer immunotherapy, early phase clinical trials have shown limited signs of clinical benefit, suggesting that blockade of CD47 alone may not be sufficient for effective tumor control. Here, we investigate the contributions of the Fc domain of anti-CD47 antibodies required for optimal in vivo antitumor activity across multiple species-matched models, providing new insights into the mechanisms underlying the efficacy of this emerging class of therapeutic antibodies. Using a novel mouse model humanized for CD47, SIRPα and FcγRs, we demonstrate that local administration of an Fc-engineered anti-CD47 antibody with enhanced binding to activating FcγRs modulates myeloid and T-cell subsets in the tumor microenvironment, resulting in improved long-term systemic antitumor immunity and minimal on-target off-tumor toxicity. Our results highlight the importance of Fc optimization in the development of effective anti-CD47 therapies and provide a novel approach for enhancing the antitumor activity of this promising immunotherapy.

**Highlights:**

- Engagement of activating FcγRs augments the in vivo antitumor activity of CD47 blocking antibodies
- Humanized mice for CD47, SIRPα and FcγRs allow assessment of hFcγRs contribution to the activity of anti-hCD47 Abs
- Fc-optimized anti-hCD47 ab promotes systemic antitumor immunity with abscopal effect and minimal on-target toxicity

**Graphical Abstract:** 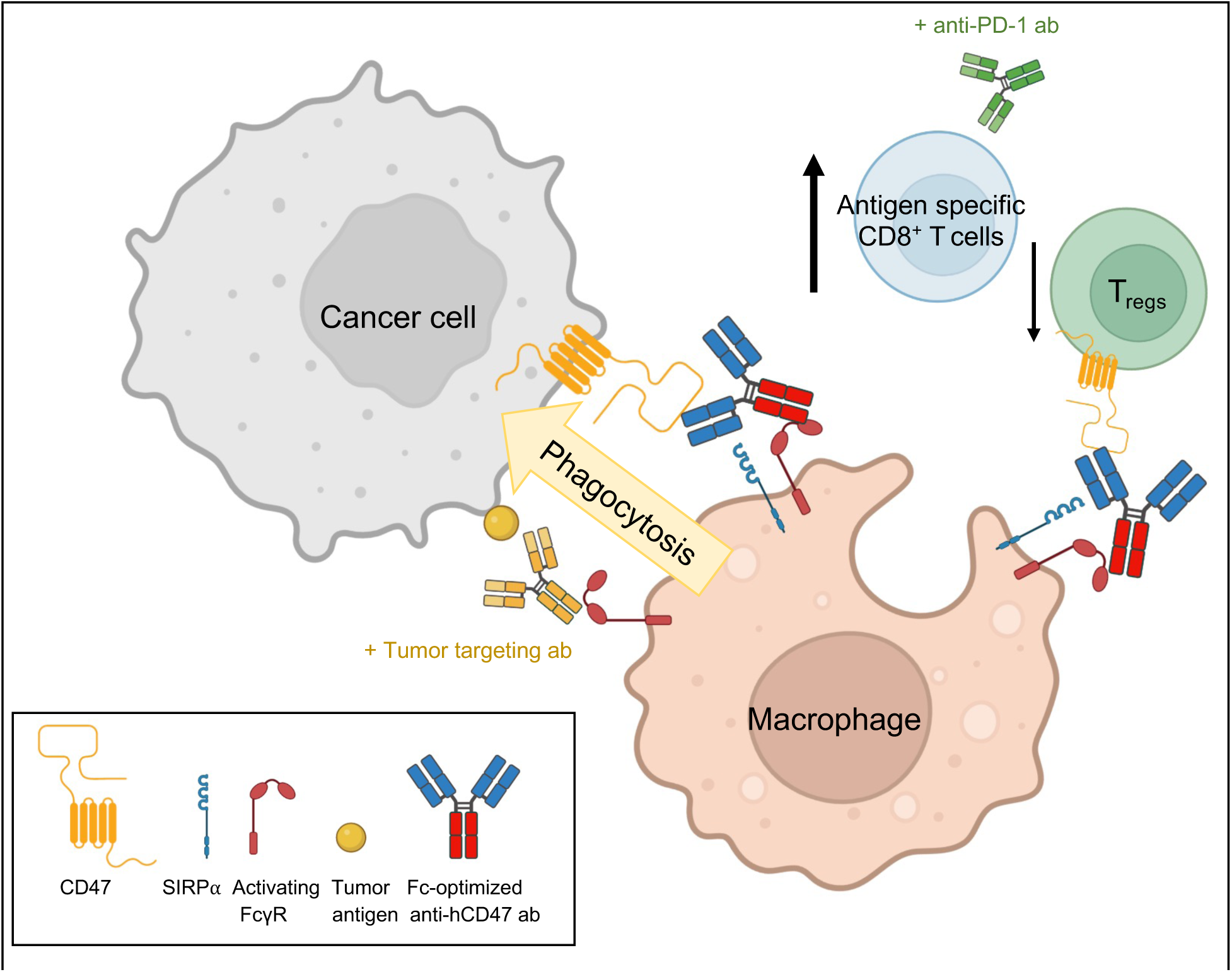

## Introduction

Antibodies (Abs) blocking inhibitory immune checkpoints expressed primarily on adaptive immune cells lead to profound and durable clinical responses in a wide range of malignancies (Ribas and Wolchok, 2018). However, the benefit of these therapies is limited only to a subset of patients. As a result, there is a growing interest in identifying additional checkpoints for cancer immunotherapy including those expressed on innate immune cells. The CD47/SIRPa axis has emerged as an essential checkpoint for cancer immunosurveillance by functioning as a “don’t eat me” signal that suppresses innate immune responses. CD47 is a cell surface receptor expressed on healthy cells and often overexpressed on cancer cells(Willingham et al., 2012), which protects them from phagocytosis by interacting with SIRPα, an inhibitory immunoreceptor expressed on myeloid cells including macrophages, neutrophils and subsets of dendritic cells(Adams et al., 1998). In several preclinical studies, therapeutic antibodies that block the interaction between CD47 and SIRPα promote effective antitumor responses by enabling elimination of both hematologic and solid tumor cells by phagocytes (Chao et al., 2010; Jaiswal et al., 2009; Majeti et al., 2009; Weiskopf et al., 2016; Willingham et al., 2012), enhancing cross-presentation of CD8^+^ T cells by antigen presenting cells, and triggering adaptive immune responses(Liu et al., 2016; Liu et al., 2015b). These encouraging results led the development of several antibodies and fusion proteins blocking CD47, which are now being investigated in over 50 clinical trials across multiple tumor types (Jalil et al., 2020; Uger and Johnson, 2020).

Despite these advances, anti-CD47 antibodies have demonstrated limited clinical benefit when administered as monotherapy(Bouwstra et al., 2022), highlighting the need to better understand the mechanisms behind the efficacy of these therapies. To this end, a critical question that remains to be answered is whether the antitumor activity of these antibodies relies solely on blocking the CD47/SIRPα interaction by the fragment antigen-binding (Fab) domain, or whether interactions between the fragment crystallizable (Fc) and Fc gamma receptors (FcγRs) also contribute to their antitumor activity(Zhao et al., 2012a; Zhao et al., 2012b). FcγRs are functionally classified as activating (FcγRI, FcγRIIA, FcγRIIIA in humans; FcγRI, FcγRIII, FcγRIV in mouse) or inhibitory (FcγRIIB in both human and mouse), and are expressed on a variety of cells (Nimmerjahn and Ravetch, 2008). The effector responses mediated by different IgG subclasses are dependent on their differential affinities for activating or inhibitory FcγRs. As a result, FcγR signaling through the Fc portion of IgG antibodies can induce activation of phagocytic and cytotoxic cells, and has been found essential for the therapeutic activity of antibodies targeting tumor antigens (Bevaart et al., 2006; Clynes et al., 2000; DiLillo and Ravetch, 2015), and most recently on antibodies blocking or agonizing immune checkpoints(Dahan et al., 2016; Dahan et al., 2015). Optimization of these interactions have led the development of second-generation antitumor immunotherapies to further enhance their therapeutic potential (Bournazos et al., 2017; Cohen Saban et al., 2023; Garris et al., 2021; Knorr et al., 2018; Liu et al., 2020).

Currently, there is no consensus on the ideal Fc format that will maximize the therapeutic index of Ab targeting CD47. This is evident by the variety of IgG subclasses used to design these therapies for clinical trials (Jalil et al., 2020; Uger and Johnson, 2020). Results from early-phase clinical trials indicate that antibodies with weak or null engagement to FcγRs (hIgG4, hIgG2 or inert Fc region) were relatively tolerable, but did not show clinical benefit when administered as monotherapy (Burris et al., 2021; Sikic et al., 2019; Zeidan et al., 2022). Of note, they only demonstrated benefit in combination with other antitumor antibodies that engage FcγRs such as Rituximab (Abrisqueta et al., 2019; Advani et al., 2018). In contrast to these results, antibodies with competent binding to FcγRs (hIgG1 Fc format) showed the highest levels of therapeutic activity as monotherapy(Ansell et al., 2021; Horwitz et al., 2021; Querfeld et al., 2018); however, low concentrations of the drug were administered due to transient depletion normal CD47-expressing cells such as red blood cells (RBC) and platelets, suggesting that the Fc could also mediate some of the on-target off-tumor toxicity. Overall, these results suggests that simple blockade of the CD47-SIRPα interaction may not be sufficient to induce significant antitumor immunity and that concomitant pro-phagocytic signals, such as engagement of FcγRs may be also necessary. Optimizing the capacity of CD47 blocking antibodies to engage appropriate FcγR pathways is expected to promote activation of effector cells in the tumor microenvironment (TME), resulting in significant improvement in their in vivo antitumor activity.

Most preclinical investigations on the therapeutic activity of anti-CD47 antibodies used xenograft(Edris et al., 2012; Weiskopf et al., 2016; Xu et al., 2015) or immune-compromised(Chao et al., 2010; Jaiswal et al., 2009; Majeti et al., 2009) mouse models. These are limited by substantial interspecies differences between mouse (m) and human (h) CD47-SIRPα as well as Fc-FcyR interactions(Kwong et al., 2014; Nimmerjahn and Ravetch, 2008; Subramanian et al., 2006), providing a potential explanation as to why they have failed to correlate with efficacy and toxicity observed in patients. In this study, we provide new insights into the mechanisms behind the therapeutic activity of antibodies targeting CD47 by elucidating the effector functions engaged by Fc-FcγR interactions in several immune competent, species-matched tumor models, including a novel mouse model humanized for CD47, SIRPα and FcγRs, which overcomes the limitations of prior studies. In this platform, we evaluate whether FcγR singling can be exploited to further augment the antitumor immunity of these therapies and allow us to identify an Fc-optimized antibody targeting human CD47 for effective in vivo antitumor activity.

## Results

### Engagement of activating FcγRs contributes to in vivo antitumor activity of anti-mCD47 antibodies

To determine the contribution of Fc-FcγR interactions to the in vivo activity of antibodies targeting the CD47-SIRPα axis, we modified the Fc domain of the anti-mCD47 antibody (ab) clone MIAP301 to generate antibodies with varying affinity to mFcγRs: 1) mIgG2a Fc, binding preferentially to activating mFcγRs I and IV, 2) mIgG1 Fc, binding preferentially to the inhibitory mFcγRIIB, and 3) mIgG1-D265A Fc, which lacks binding to any mFcγRs (**Figure S1A)** (Nimmerjahn and Ravetch, 2005). Alterations to the Fc domain of MIAP301 did not alter the binding specificity of the variable regions to mCD47 (**Figure S1C)**, nor its in vivo half-life (Dahan et al., 2015) which allows comparisons into the contributions of differential FcγR engagement to the activity of CD47 blocking antibodies. We compared the in vivo antitumor activity of these different Fc subclasses of MIAP301 in wild type B6 (WT) mice bearing subcutaneous (sq.) MC38 colorectal adenocarcinomas, a tumor type expressing high levels of CD47 **(Figure S1B)**. We found the mIgG2a Fc variant of MIAP301 led to the most significant reduction in tumor burden when compared to the control or other Fc variants **(Figure 1A and S2A).** Of note, this therapeutic effect was attenuated for the mIgG1-D265A Fc variant and in mice lacking all activating FcγRs (FcR-γ chain^-/-^) (Takai et al., 1994) **(Figure 1A and S2B).** The mIgG2a Fc variant of MIAP301 also demonstrated the most significant decrease in the number of lung metastases in a second tumor model using B16 melanoma cells **(Figure 1B and S2C).**

**Figure 1.**
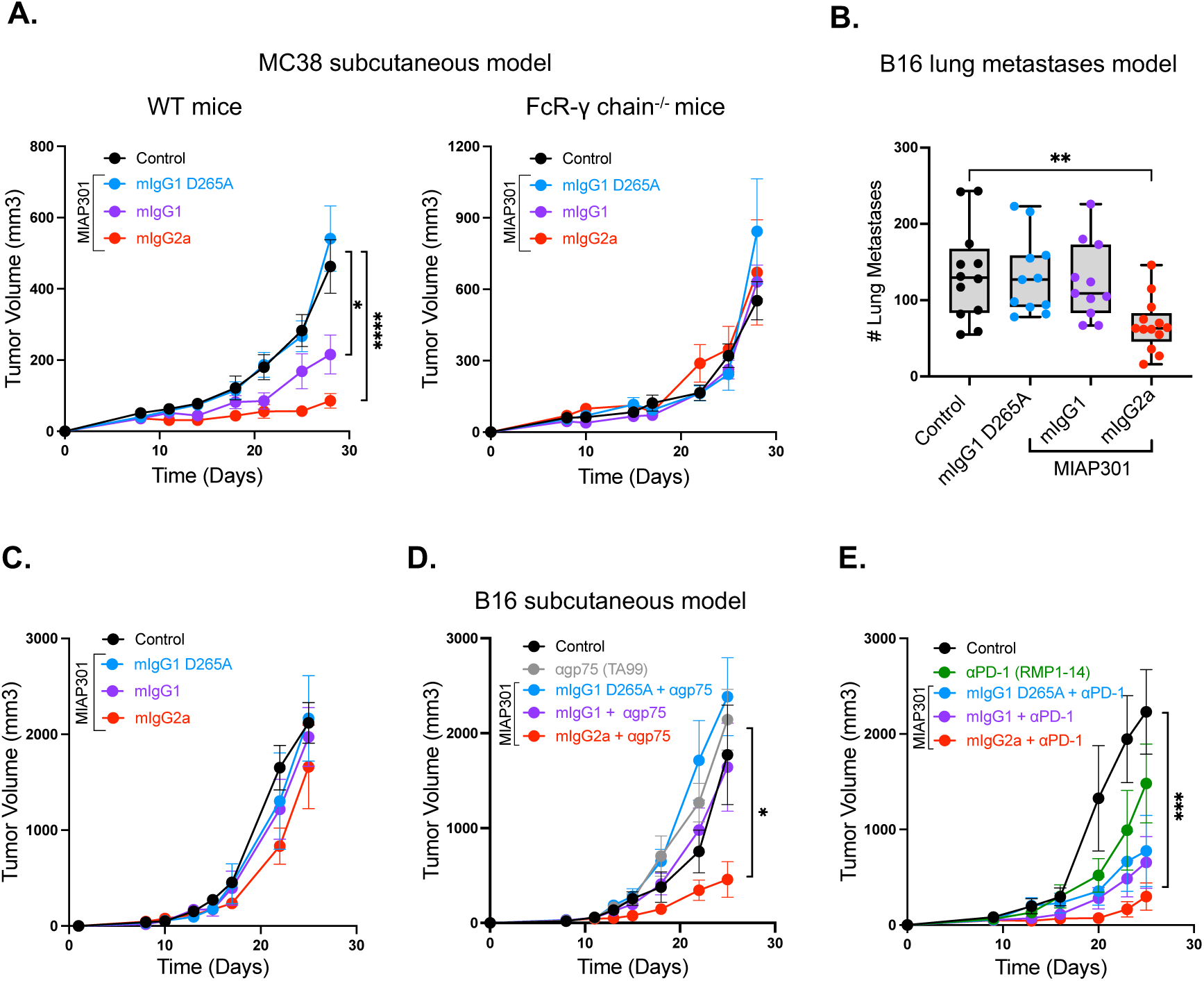
Engagement of activating FcγRs enhances in vivo antitumor activity of antibodies blocking mouse CD47. **(A)** Average growth ± SEM of sq. MC38 tumors in WT (right) or FcR-γ chain^-/-^ mice (left), treated with Fc variants of anti-mCD47 ab (MIAP301) or control (50ug IT, d. 8,10,14 and 18). **(B)** Average of lung metastases ± (Min to Max) of WT mice inoculated IV with B16 tumor cells, treated with anti-mCD47 ab Fc variants (MIAP301) or control (20 mg/Kg IP, d. 1,4,7 and 11). **(C)** Average growth ± SEM of sq. B16 tumors in WT mice, treated with MIAP301 Fc variants or control (50ug IT, d. 8,10,14 and 18). **(D)** Average growth ± SEM of sq. B16 tumors in WT mice, treated with MIAP301 Fc variants or control (50 ug IT, d. 8,10,14 and 18) in combination with anti-gp75 ab (TA99-mIgG2a, 200 ug IP d. 8, 10, 14 and 18). **(E)** Average growth ± SEM of sq B16 tumors in WT mice, treated with MIAP301 Fc variants or control (50ug IT, d. 8,10,14 and 18) in combination with anti-PD-1 ab (RMP1-14-mIgG1-D265A, 200 ug IP d. 8, 10, 14 and 18).

Currently, there are several clinical trials investigating anti-CD47 antibodies in combination with therapies blocking tumor antigens of immune checkpoints (Maute et al., 2022). We thus compared the anti-tumor activity of the different Fc variants of MIAP301 in combination with a tumor targeting ab against gp75 (Clone TA99, mIgG2a ab) (Clynes et al., 1998) or with an ab blocking the immune checkpoint PD-1 (Clone RMP1-14, mIgG1 D265A ab). To assess these combinations, we used a sq. B16 melanoma model in which none of our MIAP301 Fc variants worked as monotherapy **(Figure 1C and S2D).** The combination of anti-gp75 ab with the MIAP301-mIgG2a ab led to significant reduction in tumor burden when compared to anti-gp75 ab alone or the other combinations **(Figure 1D and S2E).** Similarly, combination of the anti-PD-1 ab with MIAP301-mIgG2a ab significantly decreased tumor burden when compared to PD-1 blockade alone or the other combinations **(Figure 1E and S2F).** The enhanced activity of the mIgG2a ab compared to the other Fc subclasses alone or in combination with tumor targeting antibodies or immune checkpoint blockade indicates that engagement of activating FcγRs by CD47 antibodies contributes to its optimal in vivo anti-tumor activity.

### Engagement of activating FcγRs during treatment with anti-CD47 antibodies correlates with increased number of tumor-infiltrating macrophages and enhanced phagocytic activity

To evaluate the factors that promote the enhanced activity of the mIgG2a Fc subclass of MIAP301, we compared the effect of the three Fc variants on myeloid cells in the TME. Treatment with the MIAP301-mIgG2a ab correlates with a significant increase in the percentage of CD45^+^ leukocytes **(Figure S3A)**, mainly attributed to a significant increase in CD11b^+^F4/80^+^ macrophages and CD11b^+^Ly6C^+^ monocytes, but not neutrophils or dendritic cells **(Figure 2A and S3D-E).** These findings suggested that these two populations could mediate the effector response during treatment with this Fc variant. To test this hypothesis, we selectively depleted tumor infiltrating macrophages or CCR2^+^ monocytes using clodronate or an anti-CCR2 ab (mIgG2a), respectively **(Figure S3F-H).** Depletion with clodronate – but not with anti-CCR2 ab – abrogated the activity of the MIAP301-mIgG2a ab **(Figure 2B-C)**, indicating that macrophages are required effector cells for the increased therapeutic activity of MIAP301-mIgG2a. Since CD47 constitutes a well-known antiphagocytic signal on macrophages, we assessed the effect of the different anti-mCD47 Fc subclasses in macrophage activity by performing in vitro phagocytosis assays of MC38 and B16 cancer cells using bone marrow derived macrophages (BMDM). The MIAP301-mIgG2a ab led to significantly more phagocytosis of both cancer cell lines when compared to control or the other Fc subclasses **(Figure 2D and S4A).** These effects were also seen when other murine cancer cells lines were used as targets **(Figure S4B).** Altogether, these results indicate that engagement of activating FcγRs by CD47 antibodies increases the number of tumor-infiltrating macrophages and enhances their tumor phagocytic activity.

**Figure 2.**
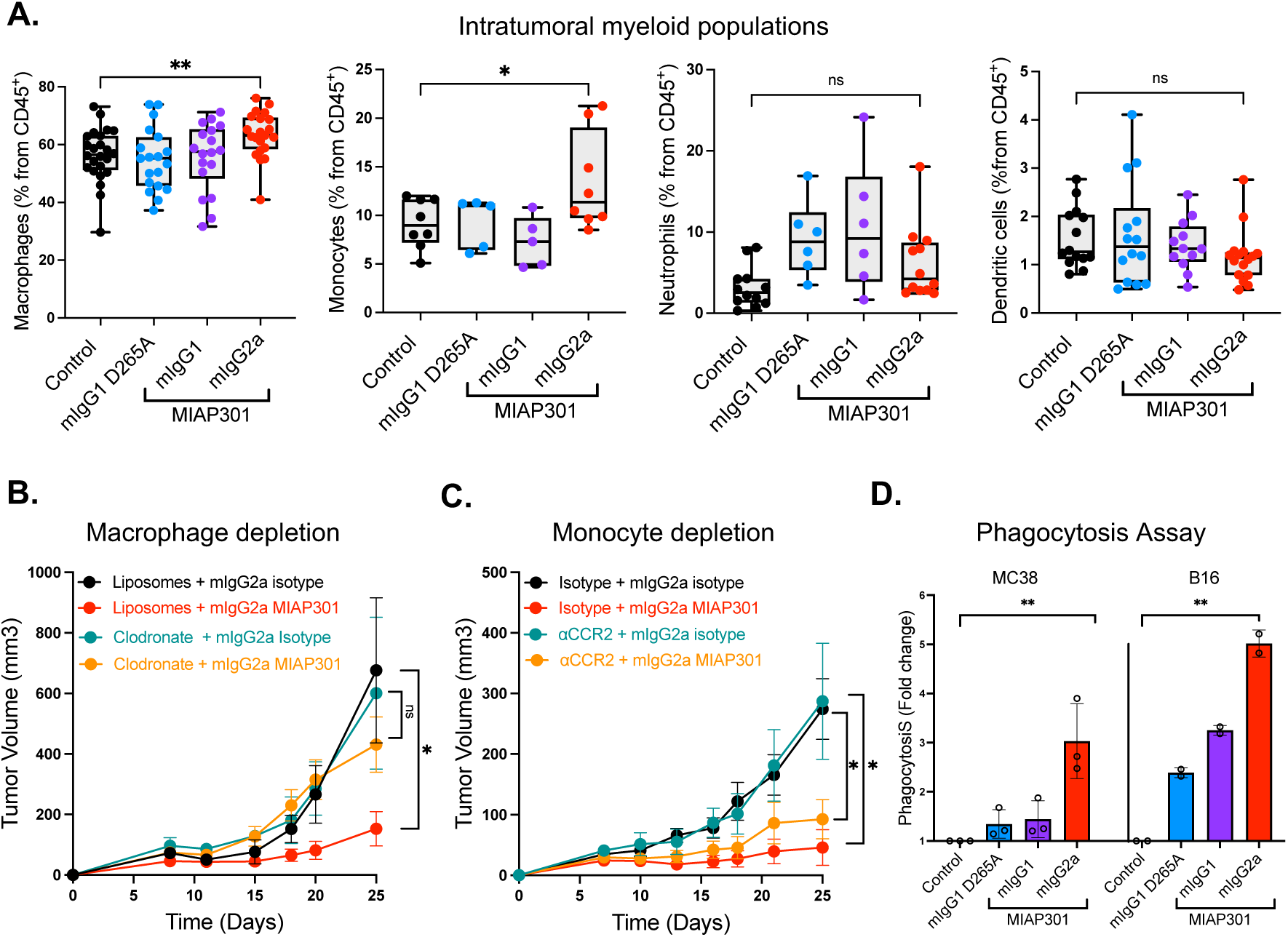
Anti-mCD47 ab Fc-enhanced for activating FcγRs correlates with increased number of tumors infiltrating macrophages and enhanced phagocytic activity. **(A)** Established MC38 tumors were analyzed by flow cytometry 72 hours after treatment with Fc variants of anti-mCD47 ab (MIAP301). Quantification of intratumoral myeloid cell populations is shown. **(B)** Average growth ± SEM of sq. MC38 tumors pre-treated with clodronate- or control-liposomes (100 uL, d.4, 7), then treatment with MIAP301-mIgG2a or isotype control was given (50 ug IT, d. 8, 10, 14 and 18). **(C)** Average growth ± SEM of subcutaneous sq. MC38 tumors pre-treated with anti-CCR2 ab (MC-21) or isotype control (100 ug, d. 7), then treatment with MIAP301-mIgG2a or isotype control was given (50 ug IT, d. 8, 10, 14 and 18). **(D)** Bone marrow derived macrophages obtained from WT mice were cocultured with CFSE-labeled MC38 or B16 tumor cells in the presence of PBS (Control) or Fc-modified anti-mCD47 ab (MIAP301). Percentage of phagocytosis was determined by the percentage of CFSE^+^ cells within the macrophage cell gate (CD11b^+^F480^+^). Analysis on fold changes compared to control is shown.

### Engagement of activating FcγRs during treatment with anti-CD47 antibodies modulates subsets of T cells and promotes long term antitumor immunity

CD47 blockade promotes adaptive immune responses by increasing cross-presentation of tumor antigens (Liu et al., 2015b; Tseng et al., 2013) and by inducing type 1 immune responses through direct ligation of CD47 expressed on T cells (Bouguermouh et al., 2008). We first analyzed the patterns of CD47 expression on CD8^+^, CD4^+^FOXP3^-^, and CD4^+^FOXP3^+^ (Regulatory T cells or T_regs_) T cells in the TME. We found that all three T cell subsets expressed CD47, however T_regs_ exhibited a significantly higher expression when compared to CD4^+^FOXP3^-^ or CD8^+^ T cells **(Figure 3A).** This differential expression of CD47 supports the possibility that the anti-tumor effect observed after treatment with an Fc-enhanced anti-CD47 ab results from both enhanced phagocytosis and depletion of T_regs_ in the TME. To test this hypothesis, we assessed the effect of the three sub-classes of MIAP301 on T_regs_ in the TME of MC38 tumors. Treatment with the MIAP301-mIgG2a ab led to a significant decrease in the percentage of T_regs_ when compared to control tumors **(Figure 3B),** suggesting that treatment with the MIAP301-mIgG2a ab is associated with an Fc-mediated decrease in the frequencies of immunosuppressive CD47^high^ expressing T_regs_.

**Figure 3.**
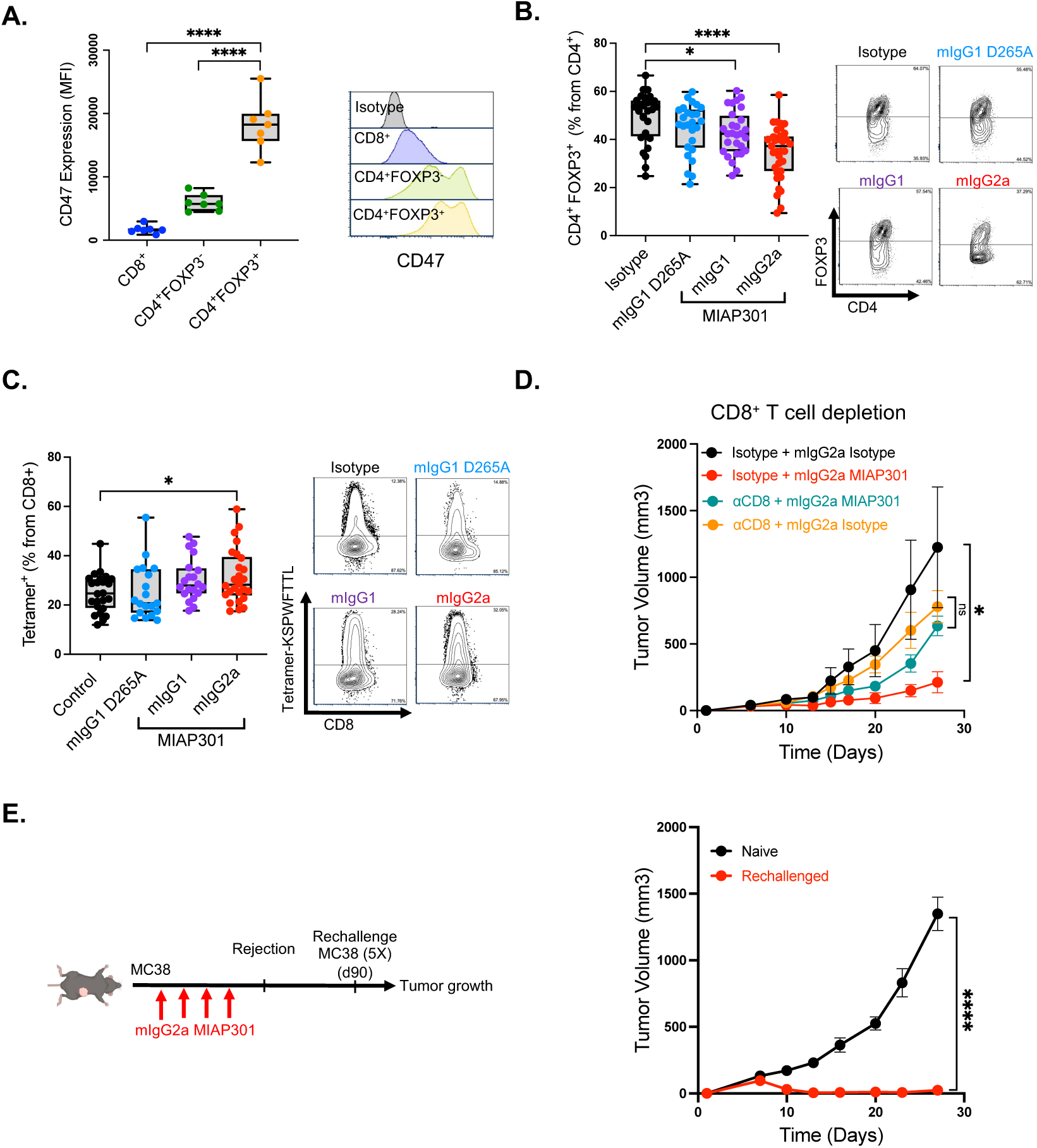
Fc-optimized anti-mCD47 antibodies decreases T_regs_ and promotes infiltration of antigen-specific CD8^+^ T cells. **(A)** Levels of expression of CD47 on T cell subsets isolated from MC38 tumors, quantification (left) and representative flow cytometry plots (right) are shown. **(B)** Percentage of CD4^+^FOXP3^+^ T_regs_ of total CD4^+^ T cells, and **(C)** Percentage of tetramer^+^ (KSPWFTTL) cells of total CD8^+^ T cells from MC38 tumors 72 hours after treatment with Fc variants of anti-mCD47 abs (MIAP301). **(D)** Average growth ± SEM of sq. MC38 tumors pre-treated with anti-CD8 ab (2.43) or isotype control (100 ug. d.7, 12, 17), then treatment with MIAP301-mIgG2a or isotype control was given (50 ug IT, d. 8, 10, 14 and 18). **(E, left)** MC38 tumors were treated with MIAP301-mIg2a (50 ug every 3 days) until rejection was achieved. Then mice were rechallenged with MC38 cells (10 million, d. 90). **(E, right)** Average growth ± SEM of sq. MC38 tumors in mice that achieved rejection 90 days before (rechallenged grouped) or in and mice without prior implantation of MC38 tumors (naïve group).

Since T_regs_ can restrain effective adaptive antitumor activity (Shan et al., 2022), we investigated whether decreased levels of T_regs_ correlate with a higher number of CD4^+^FOXP3^-^ or CD8^+^ T cells. No significant changes were seen in the percentages of these two populations, however, there was a significant increase in the number of antigen-specific T cells **(Figure 3C and S3B-C)**. Furthermore, depletion of CD8^+^ T cells abrogated the antitumor activity of the MIAP301-mIgG2a ab, indicating that CD8^+^ T cells are required for its optimal therapeutic activity **(Figure 3D)**. To determine whether tumor-specific T cell responses induced by the MIAP301-mIgG2a ab resulted in long term immune memory, WT mice in which MIAP301-mIgG2a ab led to a complete antitumor response were subsequently rechallenged with a 5-fold higher dose of MC38 tumor cells 90 days following initial tumor challenge in the absence of additional therapy **(Figure 3E).** Mice previously treated with the MIAP301-mIgG2a ab completely rejected the tumor rechallenge when compared to naïve mice **(Figure 3E)**. Overall, these findings indicate that treatment with a mIgG2a anti-CD47 ab facilitates long term CD8^+^ tumor-specific antitumor immunity.

### Generation of CD47/SIRP⍺ humanized knock-in mice to assess antibodies targeting human CD47 in vivo

Having established the contribution of mFcγR engagement in an immune competent murine system, we examined whether our findings were applicable for antibodies targeting human CD47 (hCD47). For this purpose, we developed a humanized mouse strain in which the human extracellular domains (ECD) of CD47 and SIRPα replaced their respective murine counterparts using CRISPR/Cas9 gene-editing (hCD47/hSIRPα knock in (KI) mice) **(Figure S5A).** We assessed the expression pattern of hCD47 and hSIRPα in these mice and found that, similar to the pattern seen on human cells, hCD47 is ubiquitously expressed on peripheral blood red blood cells (RBC), platelets, CD3^+^ and CD11b^+^ cells. In contrast, SIRPα was expressed primarily on CD11b^+^ cells but not in other cells **(Figure 4A).** Peripheral blood levels of RBC, platelets and other leukocytes were similar than those found WT mice **(Figure S5B).** Furthermore, the hCD47 and hSIRPα expressed on these humanized mice were capable of binding their human counterparts **(Figure S5C).** In an analogous CRISPR/Cas9 gene-editing strategy, we generated MC38 and B16 cancer cell lines in which their mouse ECD of CD47 was replaced by the hCD47 ECD (MC38-hCD47 KI, B16-hCD47 KI, respectively) **(Figure S5D-E)**. These cell lines can be engrafted in the hCD47/hSIRPα KI mouse system and represent a unique system to test anti-human antibodies.

**Figure 4.**
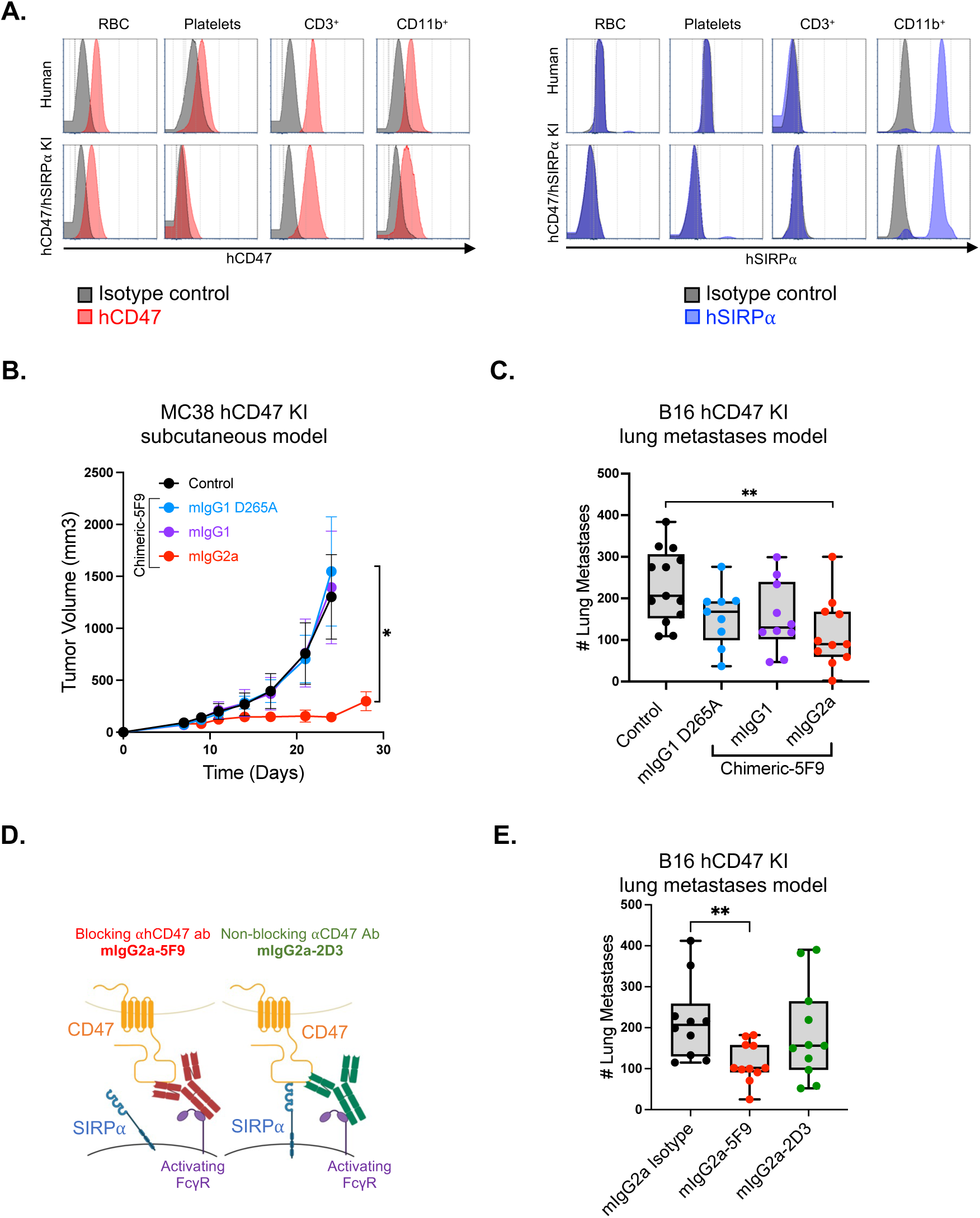
Characterization of hCD47/hSIRP⍺ KI mice and antitumor activity of blocking and non-blocking hCD47 antibodies Fc enhanced for activating FcγRs. **(A)** Flow cytometry analysis of hCD47 and hSIRP⍺ in red blood cells (RBC), platelets, CD3^+^ and CD11b^+^ leucocytes isolated from peripheral blood of human (top) and hCD47/hSIRP⍺ KI mice (bottom). **(B)** Average growth ± SEM of sq. MC38 hCD47 KI tumors in hCD47/hSIRP⍺ KI mice, treated with chimeric anti-hCD47 ab Fc variants (chimeric-5F9) or isotype control (50 ug IT, d. 8,10,14 and 18). **(C)** Average of lung metastases ± SD of hCD47/hSIRP⍺ KI mice inoculated IV with B16 hCD47 KI tumor cells, treated with chimeric anti-hCD47 ab Fc variants (chimeric-5F9) or isotype control (20 mg/Kg IP, d. 1,4,7 and 11). **(D)** Schematic drawing showing the two ab formats proposed for effective antitumor activity of anti-CD47 antibodies. Left: An ab blocking the interaction between CD47 and SIRP⍺ AND an Fc domain engaging activating FcγRs. Right: A non-blocking anti-CD47 ab AND an Fc domain engaging activating FcγRs. **(E)** Average of lung metastases ± SD of hCD47/hSIRP⍺ KI mice inoculated IV with B16 hCD47 KI tumor cells, treated with blocking (5F9-mIgG2a) and non-blocking (2D3-mIgG2a) chimeric anti-hCD47 antibodies or isotype control (20 mg/Kg IP, d. 1,4,7 and 11).

**Figure 5.**
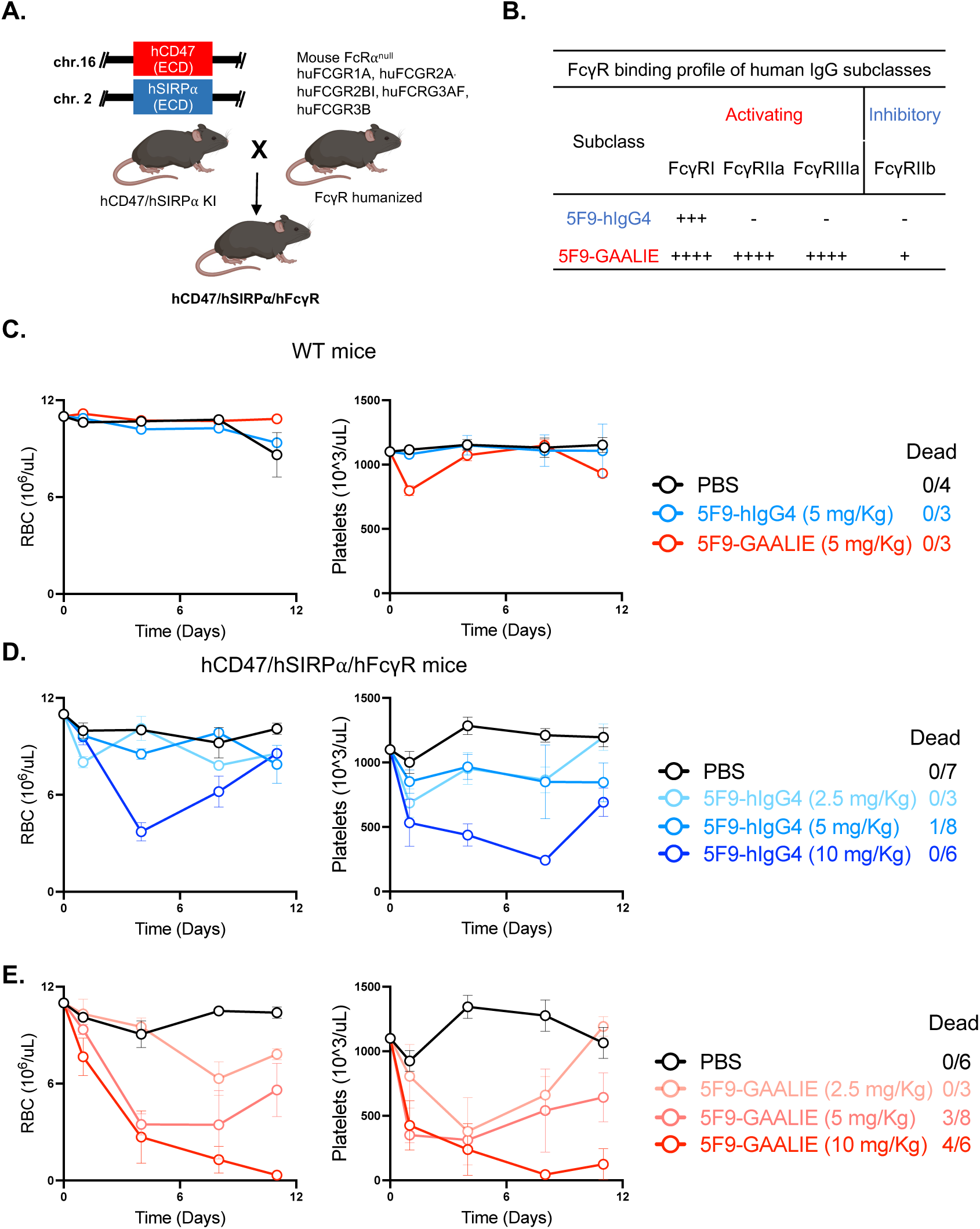
Humanized mouse model for CD47, SIRP⍺, and FcγR recapitulates activity and toxicity profile of fully humanized anti-CD47 antibodies. **(A)** Schematic drawing showing the generation of the hCD47/hSIRP⍺/hFcγR mice **(B)** Binding profiles to hFcγRs of Fc variants for anti-hCD47 abs (5F9). **(C)** Peripheral RBC and platelet counts from WT mice after treatment with 5F9-hIgG4 and 5F9-GAALIE (5 mg/Kg d. 0, 4, 8 and 11). **(D)** Peripheral RBC and platelet counts from hCD47/hSIRP⍺/hFcγR mice after treatment with increasing doses of 5F9-hIgG4 or (**E)** 5F9-GAALIE ab (2.5, 5, and 10 mg/Kg d. 0, 4, 8 and 11)

To test the therapeutic activity of antibodies against hCD47, we used the 5F9 clone (Magrolimab), which is currently being investigated in several clinical trials (Advani et al., 2018). We first generated chimeric antibodies containing human Fab and the three Fc subclasses with varying affinities to mFcγRs: 5F9-mIgG2a, 5F9-mIgG1, and 5F9-mIgG1-D265A ab **(Figure S1A and S1D)**. Using hCD47/hSIRPα KI mice, we compared the therapeutic activity of the three Fc variants of 5F9 in the MC38 hCD47KI sq. model and the B16 hCD47KI lung metastases model. Similar to the mIgG2a MIAP301 Fc variant, we found the 5F9-mIgG2a ab led to the most significant reduction in tumor burden in both tumor models when compared to control or other Fc variants **(Figure 4B-4C).** Furthermore, the 5F9-mIgG2a ab led to the most significant amount of in vitro phagocytosis of both cancer cells when compared to control or the other Fc subclasses **(Figure S4C-D).** These effects were also seen under different in vitro macrophage polarization states **(Figure S4E).** Altogether, these results validate our prior findings in the immune competent murine backgrounds and further support the role of activating FcγRs in modulating the in vivo activity of human CD47 antibodies.

### Effective in vivo antitumor activity requires both blockade of the CD47/SIRP⍺ axis and engagement of activating FcγR signaling

Using our humanized system for CD47 antibodies, we investigated whether engagement of activating FcγRs is sufficient, or whether concomitant blockade of the CD47/SIRP⍺ signal is also required to achieve optimal in vivo therapeutic activity. To address this question, we compared the in vivo antitumor activity of two antibodies targeting hCD47: The 5F9 ab clone which blocks the interaction between CD47 and SIRP⍺, and the 2D3 ab clone which binds to CD47 but does not block its interaction with SIRP⍺ (Tseng et al., 2013) **(Figure 4D).** Both antibodies were generated as mIgG2a Fc subclass to retain engagement of activating FcγRs, which allowed us to investigate the contribution of the Fab domains to the antitumor activity. We confirmed that both 5F9-mIgG2 and 2D3-mIgG2a bind to hCD47 **(Figure S1E),** but only 5F9-mIgG2a can block the interaction with hSIRP⍺ **(Figure S1F).** Furthermore, we found that similar to the 5F9-mIgG2a ab, the 2D3-mIgG2a ab is capable of inducing in vitro ADCP of cancer cells, indicating its capacity to bind FcγRs **(Figure S4F).** We compared the therapeutic activity of the two antibodies in the B16 hCD47 KI lung metastases model and found that only the 5F9-mIgG2a ab led to significantly decreased tumor burden when compared to the 2D3-mIgG2a ab or control **(Figure 4E).** These results suggest that in order to achieve significant antitumor activity, anti-CD47 antibodies require both mechanisms of activity: blockade of the CD47/SIRP⍺ (antiphagocytic) signal by the Fab domain, and engagement of the activating FcγRs (pro-phagocytic) signal by the Fc domain.

### Mice humanized for CD47, SIRP⍺ and FcγR are a reliable platform for the evaluation of toxicity and therapeutic activity of fully human anti-CD47 antibodies

After demonstrating the biological rationale for engaging activating FcγRs in both murine and human chimeric anti-CD47 antibodies, we investigated the therapeutic potential of fully humanized anti-CD47 antibodies relevant for clinical translation. Currently, most anti-hCD47 antibodies translated into patients have been studied in xenograft, immunocompromised or purely murine backgrounds(Edris et al., 2012; Liu et al., 2015b; Majeti et al., 2009; Weiskopf et al., 2016). Unfortunately, these models do not fully recapitulate with the on-target off-tumor toxicity as well as therapeutic activity observed in patients receiving these therapies as monotherapy(Sikic et al., 2019; Zeidan et al., 2022). To overcome this limitation, we crossed our hCD47/hSIRPα KI mice with a mouse humanized for FcγRs, in which all mouse FcγR genes have been deleted and all human FcγRs are expressed as transgenes (Smith et al., 2012) **(Figure 6A)**. We confirmed that our hCD47/hSIRPα/hFcγR mice maintain the human cell-type specific expression profile of FcγRs found in the humanized mice for FcγRs **(Figure S6).**

**Figure 6.**
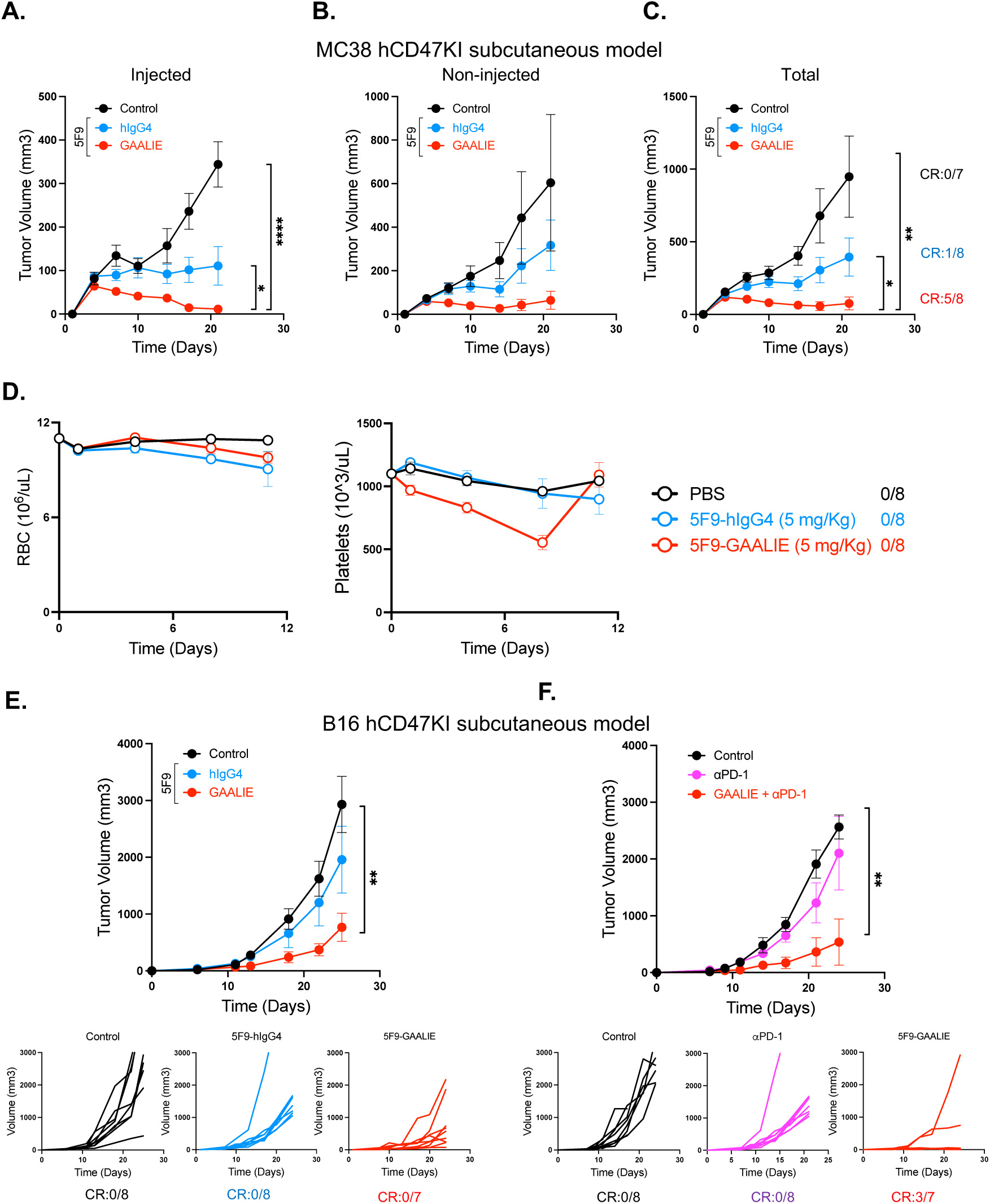
Fc-optimized humanized anti-CD47 Ab promotes effective in vivo antitumor activity. **(A)** Average growth ± SEM of injected sq. MC38 hCD47 KI tumors in hCD47/hSIRP⍺/hFcγR, treated with 5F9-hIgG4, 5F9-GAALIE ab or control (50ug IT, d. 7, 9, and 12). **(B)** Average growth ± SEM of non-injected contralateral sq. MC38 hCD47 KI tumors in hCD47/hSIRP⍺/hFcγR mice, treated with treated with 5F9-hIgG4, 5F9-GAALIE ab or control (50ug IT, d. 7, 9, and 12). **(C)** Average growth ± SEM of total (injected and non-injected) sq. MC38 hCD47 KI tumors in hCD47/hSIRP⍺/hFcγR, treated with 5F9-hIgG4, 5F9-GAALIE ab or control (50ug IT, d. 7, 9, and 12). **(D)** Serial measurements of peripheral RBC and platelets from hCD47/hSIRP⍺/hFcγR mice after treatment with 5F9-hIgG4 or 5F9-GAALIE (d. 0 indicates the day when treatment was started). **(F)** Average growth ± SEM of sq. B16 hCD47 KI tumors in hCD47/hSIRP⍺/hFcγR, treated with 5F9-hIgG4, 5F9-GAALIE ab or control (50ug IT, d. 8,10,14 and 18). **(G)** Average growth ± SEM of sq. B16 hCD47 KI tumors in hCD47/hSIRP⍺/hFcγR, treated with 5F9-hIgG4, 5F9-GAALIE ab alone (50ug IT, d. 8,10,14 and 18), or in combination with anti-PD1 ab (RMP1-14, 200 ug IP d. 8, 10, 14 and 18), or control.

First, we hypothesized that the humanized hCD47/hSIRPα/hFcγR mice could recapitulate the toxicity observed with CD47 blocking antibodies in the clinical setting, in particular anemia and thrombocytopenia(Oldenborg et al., 2000). To test this hypothesis, we compared the human CD47 blocking antibody 5F9-hIgG4 used in several clinical trials (Advani et al., 2018), with the Fc-optimized variant 5F9-GAALIE. These two antibodies have identical Fab regions, but differ in their Fc: the 5F9-hIgG4 ab is a “weak” engager of FcγRs; in contrast, the 5F9-GAALIE is a human IgG1 Fc format containing three point mutations that enhances binding for all the activating FcγRs and reduced binding to the inhibitory FcγR, thus optimizing the A/I ratio (Weitzenfeld et al., 2019) **(Figure 6B).** Systemic administration of these two human antibodies in WT mice did not lead to significant changes in peripheral RBCs or platelets in WT mice **(Figure 6C).** In contrast, increasing dosing concentration of both 5F9-hIgG4 and 5F9-GAALIE antibodies led to anemia and thrombocytopenia in hCD47/hSIRPα/hFcγR mice. Of note, these effects were transient (lowest values by day 4 to 8) and most of them recovered by day 11. While the 5F9-G4 ab started to decrease the number of RBC and platelets at a high systemic dose (10 mg/Kg) **(Figure 6D)**, the 5F9-GAALIE antibody led to anemia and thrombocytopenia at the lowest systemic dose administered (2.5 mg/Kg), with significantly more anemia, thrombocytopenia, and mortality in mice treated a higher dose (5 mg/Kg and 10 mg/Kg) **(Figure 6E).** These findings indicate that engagement of activating FcγRs during CD47 ab blockade also mediates off-tumor on-target toxicity of these therapies, and that the hCD47/hSIRPα/hFcγR mice provide a unique and reliable platform for the study of clinically relevant Fc-optimized anti-CD47/SIRPα antibodies in vivo by recapitulating important parameters of biological activity and toxicity observed in the clinical setting.

### Fc-optimized anti-CD47 antibodies promote superior antitumor activity

To evaluate the in vitro activity of the two different Fc formats of hCD47 blocking antibodies, we assessed phagocytosis of B16 hCD47 KI or human cancer cells by macrophages generated from species matched mononuclear cells. The GAALIE 5F9 ab led to significantly more phagocytosis of cancer cells when compared to the hIgG4 format or control **(Figure S4F-S4I).** Given the increased Fc-mediated toxicity observed with systemic administration of the 5F9-GAALIE ab, we next assessed whether direct (intratumoral, IT) delivery of the lowest dose (2.5 mg/kg) of anti-hCD47 antibodies to the tumor site could help control tumor growth while providing an optimal safety profile. Intratumoral administration of the 5F9-GAALIE ab led to a significant reduction in tumor burden when compared to the 5F9-IgG4 ab or control in the MC38 hCD47 KI tumor model **(Figure 6A).** Simultaneously, we evaluated whether this approach could also help control distant (non-injected) tumor growth by inducing an abscopal effect. For this purpose, we used a bilateral flank MC38 hCD47 KI tumor model which allows monitoring of tumors at both treated and untreated sites. We found that the 5F9-GAALIE ab led to significant control of distant tumor growth when compared to control, ultimately leading to complete responses in 5 out of 8 mice **(Figure 6B-C).** Of note, this local dose delivery system also resulted in significantly less toxicity **(Figure 6D).** To investigate the effects of the 5F9-GAALIE ab in a more aggressive tumor model that is traditionally resistant to single agent immune-based therapies (e.g., anti-PD-1), we used the B16 hCD47 KI sq. model (Dahan et al., 2015; Wei et al., 2017). We compared the antitumor activity of the two Fc-formats for anti-CD47 ab and found the 5F9-GAALIE ab led to a significant decrease in tumor burden as monotherapy when compared to 5F9-hIgG4 or control **(Figure 6E).** Furthermore, combination therapy with intratumoral 5F9-GAALIE ab and systemic PD-1 blockade elicited significant improved control of tumor growth when compared to PD-1 blockade administered as monotherapy or control **(Figure 6F)**. These results indicate that intratumoral dosing of Fc-optimized anti-hCD47 ab promotes effective antitumor activity, leads to abscopal immunity in distant tumor lesions and synergizes with PD-1 blockade while limiting on-target off-tumor toxicity.

## Discussion

The Fc region of antibodies blocking CD47 has been postulated as a potential modulator of their therapeutic activity (Bouwstra et al., 2022; Maute et al., 2022; Zhao et al., 2012b). However, efforts to address this question have been limited to suboptimal preclinical studies, limiting their translational relevance for human therapeutics. In this study, we overcome these limitations by elucidating the in vivo contributions of the Fc region of anti-CD47 antibodies using a novel syngeneic immunocompetent murine model that accurately reflects the affinities and cellular expression of human CD47-SIRP⍺ and IgG-FcγRs axes. As a proof of concept, we use multiple species matched systems to demonstrate that anti-CD47 antibodies displayed enhanced in vivo antitumor activity only when the antibody Fc was optimized to engage activating FcγRs. This effect depends on both macrophages and antigen specific T cells and leads to long-lived systemic immunity. Using this unique platform, we show enhanced in vivo antitumor activity, abscopal antitumor effect and minimal on-target toxicity induced by local administration of the fully human Fc-optimized anti-CD47 ab 5F9-GAALIE as monotherapy or in combination with PD-1 blockade. These data support the importance of Fc optimization in the development of effective anti-CD47 therapies and provide a reliable platform to optimize the therapeutic activity of this promising immunotherapy.

Our studies demonstrate that anti-CD47 antibodies require two mechanisms of action for optimal in vivo activity: 1) Fab-mediated CD47/SIRPα-blocking signal; and 2) Fc-mediated engagement of activating – but not inhibitory – FcγRs. This dual mechanism of action has been considered a critical factor for the activity of other immunomodulatory antibodies. This is the case of antibodies blocking CTLA-4, CCR8, OX-40, 401BB or PD-L1, in which engagement of activating FcγRs seems to augment their therapeutic activity(Arce Vargas et al., 2018; Buchan et al., 2018; Bulliard et al., 2014; Cohen Saban et al., 2023; Knorr et al., 2023; Simpson et al., 2013). In contrast, selective engagement of the inhibitory FcγRIIB enhances the activity of agonistic antibodies targeting CD40(Li and Ravetch, 2011; White et al., 2011). In certain cases, engagement of FcγRs could also compromise the activity of these therapies, which is the case of anti-PD-1 antibodies(Dahan et al., 2015). Furthermore, we observed that optimization of the Fc region of the fully humanized anti-CD47 antibody to selectively engage activating FcγRs significantly augmented the therapeutic activity of these agents in a system that closely recapitulate both human CD47-SIRP⍺ and IgG-FcγR signaling. This is important, given the extensive differences on the pattern of cellular expression and distribution between the mouse and human FcγRs of macrophages and other immune cells in the TME(Bournazos, 2019). These differences further underscores the advantage of using fully humanized Fc-FcγR systems to better estimate the activity of these therapies in humans(Bournazos et al., 2014). Altogether, we propose the clinical development and evaluation of second-generation CD47 blocking antibodies with enhanced Fc effector function, which could improve their therapeutic activity and overcome some of the limited clinical benefit observed in early-phase clinical trials.

The improved in vivo activity of anti-CD47 antibodies Fc-enhanced for activating FcγRs is characterized by modulation of several immune cell populations in the TME. Notably, we found a significant increase in macrophages in the TME of treated mice, which likely contribute to their antitumor activity. The central role of macrophages on the in vivo Fc-mediated activity of anti-CD47 antibodies is consistent with our findings and others indicating that Fc-FcγR interactions are required to induce significant phagocytosis of cancer cells by human macrophages during treatment with CD47 therapies (Jain et al., 2019; Metayer et al., 2017). This suggest that in order to achieve effective antitumor activity, anti-CD47 antibodies must not only disrupt CD47-SIRP⍺ interactions in macrophages but also opsonize the tumor cells through antibody-dependent cellular phagocytosis (ADCP). Furthermore, downstream interactions between SIRPα and FcγR signaling seem to be critical for the effector activity of these cells, with evidence indicating the ratio of activating IgG to inhibitory CD47 blockade dictates macrophage phagocytic activity by increasing phosphorylation of ITAMs on FcγRs(Suter et al., 2021).

In addition to the increased number of macrophages in the TME, we observed a decrease in the frequency of T_regs_, and an increase in the number of antigen-specific T cells. T_regs_ are a specialized sub-population of T cells that diminish effective antitumor immune responses and are an attractive target for cancer immunotherapy (Togashi et al., 2019). Of note, we observed that T_regs_ express high levels of CD47 when compared to other CD4^+^ or CD8^+^ T cells, a finding that has been observed in CTLA-4^+^ T_regs_ obtained from human tumor samples(Zhang et al., 2021). While reduced differentiation or proliferation of T_regs_ could be induced by “reprogrammed” macrophages after Fc-mediated phagocytosis of cancer cells(Tseng et al., 2013), our data suggest that Fc-optimized anti-CD47 antibodies promote T_reg_ depletion likely contributing to a less immunosuppressive TME. Nevertheless, our findings suggest that in addition to modulation of myeloid effector activity, Fc-optimized anti-CD47 antibodies also promote systemic and long-lasting adaptive immune responses by downregulating immunosuppressive T cell populations and enabling effective tumor antigen-specific cytotoxic T cell responses. The alterations in the frequency of these T cell populations in the TME may also explain the abscopal effects observed after local administration of the CD47 antibody 5F9-GAALIE, as well the long-term immune memory in the tumor rechallenge experiments.

On-target toxicity resulting from phagocytosis of normal CD47-expressing cells (i.e., RBCs and platelets) during treatment with anti-CD47 antibodies remains a concern for the clinical implementation of these therapies. The lack of cross-reactivity between most human CD47 blocking antibodies and their murine counterparts has been a major limitation of murine models. (Huang et al., 2017). Furthermore, these systems may overestimate the efficacy of these therapies as they failed to account for the large CD47 antigenic sink from normal cells. Nonhuman primates are a better system to assess toxicity(Liu et al., 2015a; Meng et al., 2019; Puro et al., 2020), however their use for simultaneous assessment of antitumor activity or alternative delivery or dosing systems are limited. Our humanized model overcomes these limitations and provides a system for simultaneous assessment of on-target off-tumor toxicity and antitumor activity. Our results indicate that the Fc region of CD47 antibodies contributes to such toxicities. These results have significant implications on the development of novel CD47 blocking agents that balance the anti-cancer Fc-mediated effects with the potential hazardous off-tumor side effects. To overcome this limitation, we tested intratumoral delivery of the 5F9-GAALIE ab, which maximizes durable control of local and distant tumor growth while minimizing on-target toxicity. Finally, we observed that combination with therapies, such as PD-1 blockade, can further enhance the systemic long-term immunity and is an attractive potential avenue to explore in patients.

The CD47-SIRPα axis is a promising target in cancer immunotherapy with a rapid developing field in recent years. By elucidating the in vivo contributions of FcγR signaling in the antitumor activity and toxicity of fully humanized anti-CD47 antibodies, our findings address a critical question in the field that will inform the rational development of Fc-optimized anti-CD47 antibodies.

## Methods

### Mouse strains

C57BL/6J (WT) (Strain#: 000664) mice were purchased from The Jackson Laboratory. Mice humanized for FcγRs (hFcγRs mouse: murine α chain KO, FcγRI^+^, FcγRIIA^R131+^, FcγRIIB^+^, FcγRIIIA^F158+^, FcγRIIIB^+^) were generated and extensively characterized in our laboratory as previously described(Smith et al., 2012). The Fc receptor common γ-chain deficient mice (FcR-γ chain^-/-^) were generated in our laboratory as previously described (Takai et al., 1994).

Mice humanized for CD47 and SIRPα (hCD47/hSIRPα knock-in) were generated independently on the C57BL/6J background by CRISPR/Cas9-mediated gene-targeting strategy. For each gene, the extracellular domain of mouse CD47 (exon 2) and mouse SIRPα (exon 2) were excised by sgRNAs and replaced with their respective human exon 2 transgene sequences (via synthesized MegaMers, IDT) through non-homologous end joining (NHEJ). To target the mouse *CD47* and *SIRPα* genes, CRISPR sgRNAs (UpCD47C-TTGCATCGTCCGTAATGTGG**AGG,** DnCD47D-ATAGAGCTGAAAAACCGCAC**GGG** and UpSIRPαA-AAATCAGTGTCTGTTGCTGC**TGG**, UpSIRPαE-GGAACAGAGGTCTATGTACT**CGG** (PAMS in bold) were used to guide the Alt-R™ S.p. HiFi Cas9 Nuclease (IDT) creating double stranded breaks.

To create the humanized CD47 knock-in, the corresponding CRISPR/Cas9 reagents were microinjected into C57BL/6 (Jackson Laboratories) 1-cell stage mouse embryos and then implanted into surrogate CD-1 mice (Charles River Laboratories). Pups born were screened for presence of the targeted allele and analyzed for proper expression. The transgenesis of the humanized SIRPα knock-in utilized a different approach to deliver the CRISPR/Cas9 kit called “improved-Genome editing via Oviduct Nucleic Acids Delivery (i-Gonad)(Ohtsuka et al., 2018). Similarly, pups born from recipient female mice were screened for presence of the Exon 2 knock-in and analyzed for proper expression. Mice humanized for hCD47 and hSIRPα (hCD47/hSIRPα KI) were generated by backcrossing a hCD47 knock-in mouse to a hSIRPα knock-in mouse. Mice humanized for CD47, SIRPα and FcγRs (hCD47/hSIRPα/hFcγR) were generated by backcrossing a fully hCD47/hSIRPα KI mouse to hFcγR mice. All mice were bred and housed at Rockefeller University Comparative Bioscience Center under specific pathogen-free conditions. All mice used were 7 to 12-week-old at time of the experiment, with a mix of male and female mice. All experiments were performed in compliance with institutional guidelines and were approved by the Rockefeller University Institutional Animal Care and Use Committee (IACUC).

### Cell lines and Cell culture

Murine MC38 colorectal cancer, B16 melanoma, Lewis Lung carcinoma (LLC) and human Jurkat and Raji lymphoma cells and were obtained from ATCC. HKP1 (Kras^G12D^p53^-/-^) lung adenocarcinoma cells were characterized and obtained from the laboratory of Dr. Vivek Mittal(Markowitz et al., 2018). All murine cells were maintained in DMEM (Life Technologies) supplemented with 10% fetal bovine serum (Life Technologies), 100 U/mL of penicillin, and 100 μg/mL of streptomycin (Life Technologies). Jurkat and Raji cells were maintained in RPMI-1640 Medium supplemented with 10% fetal bovine serum (Life Technologies, and 100 U/mL of penicillin, and 100 μg/mL of streptomycin (Life Technologies). Cells were split twice per week and cell viability was measured using trypan blue staining in the Countess II automated cell counter (Thermo Fisher). MC38 hCD47 KI and B16 hCD47 KI cancer cells were generated through CRISPR/Cas9-mediated gene targeting by replacing the extracellular domain of mouse CD47 (exon 2) by their respective human counterpart through non-homologous end joining. CRISPR sgRNA (U*C*U*GCAUAUAUGAUUAUCUU and A*C*C*UUGCAGAAGUCACUAG) were synthesized and purchased from Synthego. hCD47 (exon 2) HRD was purchased from Integrated DNA technologies (IDT). Cells were subsequently sorted using anti-hCD47 and anti-mCD47 antibodies, to obtain cells that homogeneously expressed only human CD47.

### Antibody engineering and expression

The variable heavy and light regions of anti-hCD47 ab clone 5F9 was synthesized (IDT) based on its published sequence (patent US9382320B2). The variable heavy and light regions of the anti-CD47 antibodies MIAP301 (anti-mCD47 ab) and 2D3 (nonblocking anti-hCD47 antibody) were decoded by mass spectrometry analysis (Bioinformatics Solutions, Inc.). The variable region sequences of the parental antibodies were subcloned and inserted into mammalian expression vectors with human IgG1, human IgG4, mouse IgG2, mouse IgG1 heavy chains, or human κ or mouse κ light chains, as previously described(Li and Ravetch, 2011). For the generation of Fc-domain variants of human IgG1 (GAALIE: G236A/A330L/I332E) and mouse IgG1 (D265A), site-directed mutagenesis using specific primers was performed based on the QuikChange site-directed mutagenesis Kit II (Agilent Technologies) according to the manufacturer’s instructions. Mutated plasmid sequences were validated by direct sequencing (Genewiz).

Antibodies were generated by transient cotransfection of Expi293F cells with heavy-chain and light-chain constructs. Expi293F cells were maintained in serum-free Expi293 Expression Medium and transfected using an ExpiFectamine 293 Transfection Kit (all from Thermo Fischer Scientific). Supernatants were collected 7 days after transection, centrifuged, and filtered (0.22 μm). Antibodies were purified from clarified supernatants using Protein G Sepharose 4 Fast Flow (GE Healthcare), dialyzed in PBS, and sterile filtered (0.22 μm) as previously described(Nimmerjahn et al., 2005).

### ELISA assays

Binding specificity of antibodies targeting mouse (MIAP301) and human (2D3 and 5F9) monoclonal antibodies were determined by ELISA using recombinant mouse (SinoBiological 57231-MNAH) and human (SinoBiological 12283-HCCH) CD47, respectively. Blocking activity of 5F9-mIgG2a and 2D3-mIgG2a was determined by competitive ELISA using recombinant human SIRPα-hFc (SinoBiological 11612-H02H1). The 96 well ELISA Half Area High Binding plates (Greiner Bio-One, #675061) were coated overnight at 4°C with recombinant CD47 proteins (1µg/mL). All sequential steps were performed at room temperature. After washing, the plates were blocked for 1 hour with 1X PBS containing 2% BSA), and were subsequently incubated for 1 hour with serially diluted IgGs (dilutions are indicated in the figures and were prepared in blocking solution). After washing, plates were incubated for 1 hour with horseradish peroxidase conjugated anti-human or anti-mouse IgG (Jackson IummunoResearch). For competitive ELISAs, after blocking nonspecific sites, plates were incubated for 1 hour with 1µg/mL of human SIRPα-hFc in 1XPBS with 1% BSA. After washing, plates were incubated for 1 hour with serially diluted 5F9-mIgG2a or 2D3-mIgG2a in 1X PBS with 1%BSA. After washing, plates were incubated for 1 hour with horseradish peroxidase conjugated anti-human IgG (#109-035-088, Jackson IummunoResearch). Detection was performed using a one component substrate solution (TMB), and reactions stopped with the addition of 2 M phosphoric acid. Absorbance at 450 nm was immediately recorded using a SpectraMax Plus spectrophotometer (Molecular Devices), and background absorbance from negative control samples was subtracted.

### Tumor challenge, antibody treatments, and myeloid cell depletion

MC38 (2×10^6^ cell/mouse), MC38-hCD47KI (5×10^6^ cell/mouse), B16 (5×10^5^ cell/mouse), or B16-hCD47KI (5×10^5^ cell/mouse) were inoculated subcutaneously and tumor volumes were measured biweekly with an electronic caliper and reported as volume (mm^3^) using the formula (L1^2^ × L2)/2, where L1 is the shortest diameter and L2 is the longest diameter. Mice were randomized and treated with systemic or intratumoral injections as indicated in the figures. For the lung metastases model, B16 or B16-hCD47KI (5×10^5^ cell/mouse) were inoculated IV into the lateral tail vein in 200 μL PBS. Mice were randomized and received intraperitoneal injections of 20 mg/Kg on days 1, 4, 7, and 11 after inoculation. The lungs were harvested on day 14 and analyzed for the presence of surface metastatic foci using a dissecting microscope.

Macrophage depletion was performed by intratumoral administration of clodronate liposomes (100 uL/mouse; Catalog F70101C-2, Formumax Scientific Inc) or control liposomes on days 4 and 7 after tumor cell inoculation. CCR2^+^ monocyte depletion was performed by systemic administration of anti-CCR2 antibody (Clone MC-21, 100 ug) or isotype control on day 7 after tumor inoculation. CD8^+^ T cell depletion was performed by systemic administration of anti-CD8 antibody (Clone 2.43, 100 ug, BioXcell) or isotype control on day 7 after tumor inoculation.

### Tissue processing

Tumors were dissected and cut into small pieces and transferred to gentleMACS C tubes (Miltenyi Biotec) containing enzyme mix for tough tumors (Catalog 130-096-730m Miltenyi Biotec) in Dulbecco’s modified Eagle’s medium (DMEM) (Biological Industries). Tumors were then dissociated using the gentleMACS OctoDissociator with Heaters (gentleMACS Program 37Cm_TDK_2, Miltenyi Biotec). Cell suspensions were then dispersed through a 70-μm nylon cell strainer and washed with PBS.

### Hematological analysis

50-100 μL blood samples were obtained from the retro-orbital sinus of mice under isoflurane anesthesia. Blood was placed in BD Microtainer tubes coated with ethylene diaminetetraacetic acid (K2-EDTA) and peripheral blood cell (RBC and platelets) counts were measured by Element HT5 hematology analyzer (Heska).

### Flow cytometry

Surface expression of mouse or human CD47 on murine tumor cell lines was assessed using MIAP301-PE (Biolegend) and CC2C6-APC (Biolegend) or 5F9-APC (generated in our lab) antibodies, respectively. Tumor cells (5 × 10^5^) were incubated with 0.5 μg antibody. Baseline staining was obtained using isotype-matched antibodies as controls.

Cell suspensions from tumors were isolated as described above, then stained for viability using the Aqua Amine fixable live dead dye (Thermo Fisher Scientific) in PBS at room temperature using standard protocols, then cells were washed once, and resuspended in FACS buffer (PBS with 0.5% BSA and 2 mM EDTA) with Fc blocked using human TrueStain FcX (Biolegend) and incubated in the dark for 10 min at room temperature. For cell surface staining, cells were stained in 100 uL of FACS buffer and incubated for 30 minutes at 4C. For antigen-specific T cells CD8^+^ T cells, surface staining with an MC38 tumor antigen-derived peptide KSPWFTTL-H-2K^b^ tetramer was performed (KSPWFTTL-APC, MBL)(Lee et al., 2020) For intracellular Foxp3 staining, and additional step was performed using the Foxp3/Transcription Factor Staining Bugger Set (cat. no. 00-5523, eBioscience^TM^) and FOXP3-BV421 or FOXP3-PE (Clone 150D, Biolegend) according to manufacture instructions. Cell populations were defined by the following markers: macrophages: CD45^+^, NK1.1^−^, Ly6C^−^, Ly6G^−^, CD11b^+^, F4/80^+^; monocytes: CD45^+^, NK1.1^−^, CD11b^+^, Ly6C^+^, Ly6G^−^, F4/80^−^; neutrophils: CD45^+^, NK1.1^−^, CD11b^+^, Ly6G^+^, Ly6C^low/-^, F4/80^−^; DCs: CD45^+^, NK1.1^−^, F4/80^−^, CD11c^+^, MHC-II^+^; NK cells: CD45^+^, NK1.1^+^; CD8 T cells: CD45^+^, CD3^+^, CD8^+^, CD4^−^; Tetramer^+^ CD8 T cells: CD45+, CD3^+^, CD8^+^, KSPWFTTL-H-2Kb^+^, CD4^−^; CD4 T cells: CD45^+^, CD3^+^, CD4^+^, CD8^−^, Foxp3^−^; T_regs_: CD45^+^, CD3^+^, CD4^+^, CD8^−^, Foxp3^+^. Samples were analyzed using an Attune NxT flow cytometer (Thermo Fisher) and data was analyzed using FCS Express 7 Research.

### In vitro antibody-dependent macrophage phagocytosis (ADCP)

Bone marrow cells from the tibia and femurs of B6 mice were flushed using a syringe into DMEM (Life Technologies) supplemented with 10% fetal bovine serum (Life Technologies), 100 U/mL of penicillin, and 100 μg/mL of streptomycin (Life Technologies). Cells were centrifuged followed by RBC lysis (Biolegend) for 5 min, quenched with complete media, and filtered through a 70-μM cell strainer. Cells were centrifugated and resuspended in media containing 10 ng/mL macrophage-colony stimulating factor (M-CSF) (Peprotech), and plated on 10-cm untreated petri dishes per mouse in 15 mL of media and cultured for 7 days without replenishing or changing media to derive BMDMs. For polarization to M1 macrophages, cells were washed with complete media on day 8 and were cultured in 15 mL of new media with 10 ng/mL of M-CSF, 90 ng/mL of IFN-γ (Peprotech) and 10 ng/mL of LPS for 24 hours. For polarization to M2 macrophages, cells were washed with complete media on day 8 and were cultured in 15 mL of new media with 10 ng/mL of M-CSF, 90 ng/mL of IL-4 (Peprotech) and 400 ng/mL of IL-13 (Peprotech) for 24 hours.

To quantify ADCP, tumor cells were stained with carboxyfluorescein succinimidyl ester (CFSE), washed with serum-free DMEM, and plated at a density of 150,000 cells per well in 150 μL serum-free DMEM in a 96-well ultra-low attachment round-bottom plate (Cat. no. 7007; Costar) on ice. Tumor cells were opsonized by addition of 10 μL/mL of the various Fc variants of CD47 antibodies or control for 30 min on ice. Macrophages (BMDMs or human macrophages derived from PBMCs) were harvested by cell scraping, pelleted, washed in serum-free DMEM, and added to opsonized tumor cells at a density of 50,000 cells per well in 50 μL of media for a final assay volume of 200 μL and an effector-to-tumor cell ratio of 1:3. Cells were incubated at 37 °C for 2 h, pelleted, washed FACS buffer and stained with a 1:100 dilution of anti-mF4/80 or anti-m/hCD11b antibody (Biolegend) FACS buffer for 30 min at 4 °C. Cells were pelleted, washed with TrypLE and FACS buffer, and analyzed by flow cytometry using an Attune NxT flow cytometer (Thermo Fisher) and data was analyzed using FCS Express 7 Research. Phagocytosis was determined by the percentage of CFSE^+^ cells within the macrophage cell gate (CD11b^+^/F480^+^). Results are shown as fold change when compared to the level of phagocytosis in cells treated with control.

### Statistics

Data was analyzed using Prism GraphPad software. One-way ANOVA with Tukey’s multiple comparison test was used to compare 3 or more groups. Unpaired 2-tailed *t* test was used when two groups were compared. All data, unless otherwise indicated, are plotted as mean ± standard deviation (SD). For statistical test, *P* values of ≤ 0.05 were considered to be statistically significant, indicated as \**P* ≤ 0.05, \*\**P* ≤ 0.01, \*\*\**P* ≤ 0.001, and \*\*\*\**P* ≤ 0.0001, not significant values are denoted as n.s. Lines associated with asterisks indicate groups compared for significance.

## Abbreviations

(ab): Antibody
(BMDM): bone marrow derived macrophages
(d): days
(ECD): extracellular domain
(FcγR): Fc gamma receptor
(h): human
(ITAMs): immunoreceptor tyrosine-based activation motifs
(IP): intraperitoneal
(IT): intratumoral
(KI): knock in
(m): mouse
(RBC): red blood cells
(SEM): standard error mean
(sq.): subcutaneous
(TME): tumor microenvironment
(WT): wild type.

## Acknowledgements

We thank Maria L. Baez, Alessandra E. Marino, and Carlo M. Sevilla for their excellent technical assistance. We also thank all the members of the J.V.R. Laboratory of Molecular Genetics and Immunology for helping discussions and sharing experiment materials, in particular Stylianos Bournazos for reviewing the manuscript and providing valuable feedback. M. Mack (University Hospital Regensburg) for providing us the anti-CCR2 antibody (clone MC-21)

## Author contributions

**Conceptualization,** J.C.O, D.A.K., and J.V.R.; **Methodology,** J.C.O, and D.A.K.; **Validation,** J.C.O, P.A.S., **Formal Analysis,** J.C.O, P.A.S.; **Writing,** J.C.O, D.A.K., and J.V.R.; **Resources,** J.C.O, and J.V.R.; **Supervision,** J.C.O, D.A.K., and J.V.R.

---

**Supplementary Figure 1.**
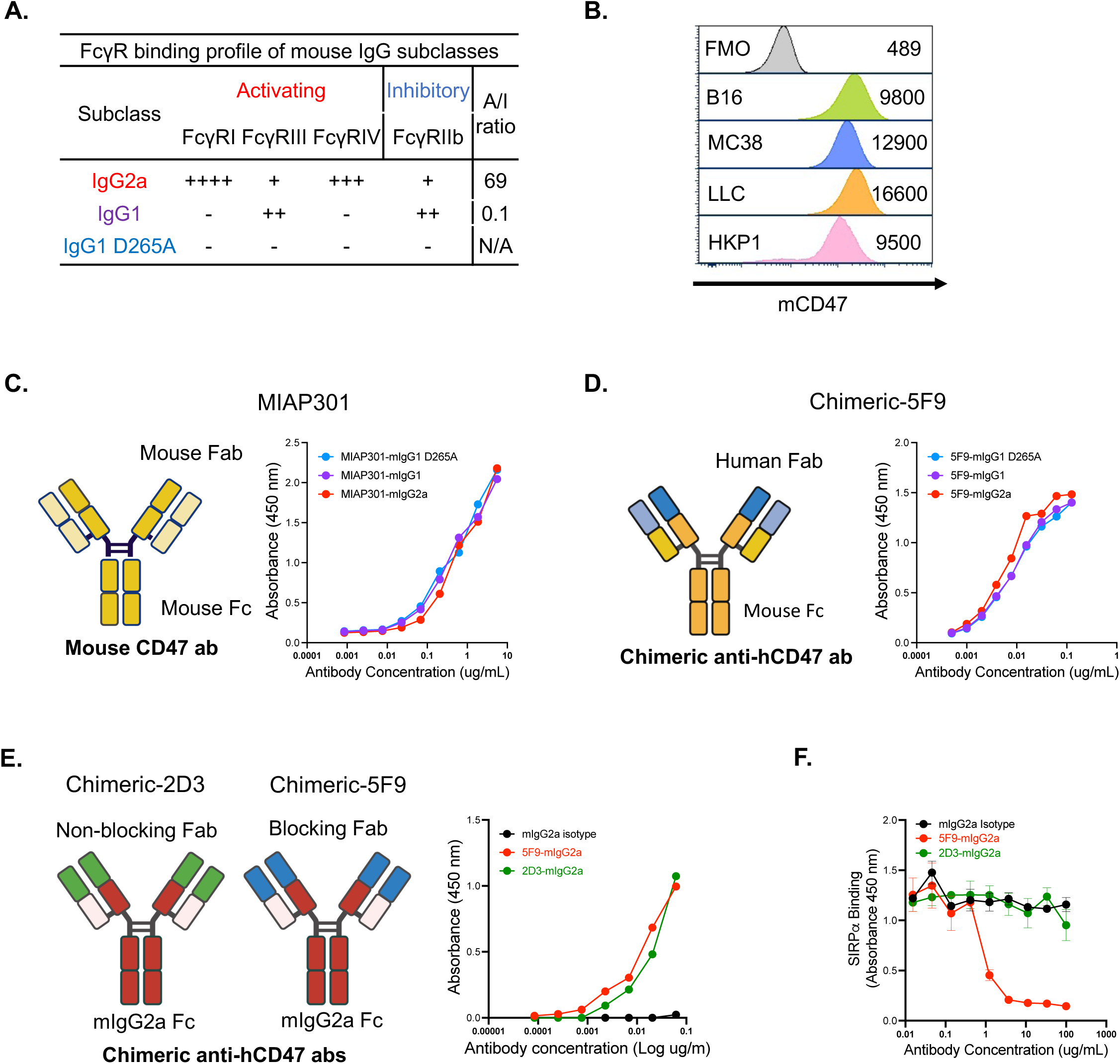
Generation and validation of Fc variants for mouse and chimeric anti-CD47 antibodies. **(A)** Binding profiles to mouse FcγRs of Fc variants for anti-CD47 antibodies (MIAP301, chimeric-5F9 and chimeric-2D3), activating-to-inhibitory binding ratio (A/I) to mFcγRs is shown **(B)** Levels of expression of CD47 on different mouse cell lines, mean fluorescence intensity (MFI) expressed on the right. **(C)** Diagram showing the structure of the mouse antibody MIAP301 and binding of their Fc variants to recombinant mouse CD47 by ELISA **(D)** Diagram showing the structure of the chimeric antibody chimeric-5F9 and binding of their Fc variants to recombinant human CD47 by ELISA **(E)** Diagram showing the structure of non-bocking (Chimeric 2D3-mIgG2a) and blocking (Chimeric 5F9-mIgG2a) CD47 antibodies. Binding to recombinant human CD47 by ELISA **(F)** Competitive ELISA showing the capacity of blocking (mIgG2a Chimeric-5F9) and non-blocking (mIgG2a Chimeric-2D3) antibodies to disrupt the binding of recombinant SIRP⍺ to hCD47.

**Supplementary Figure 2.**
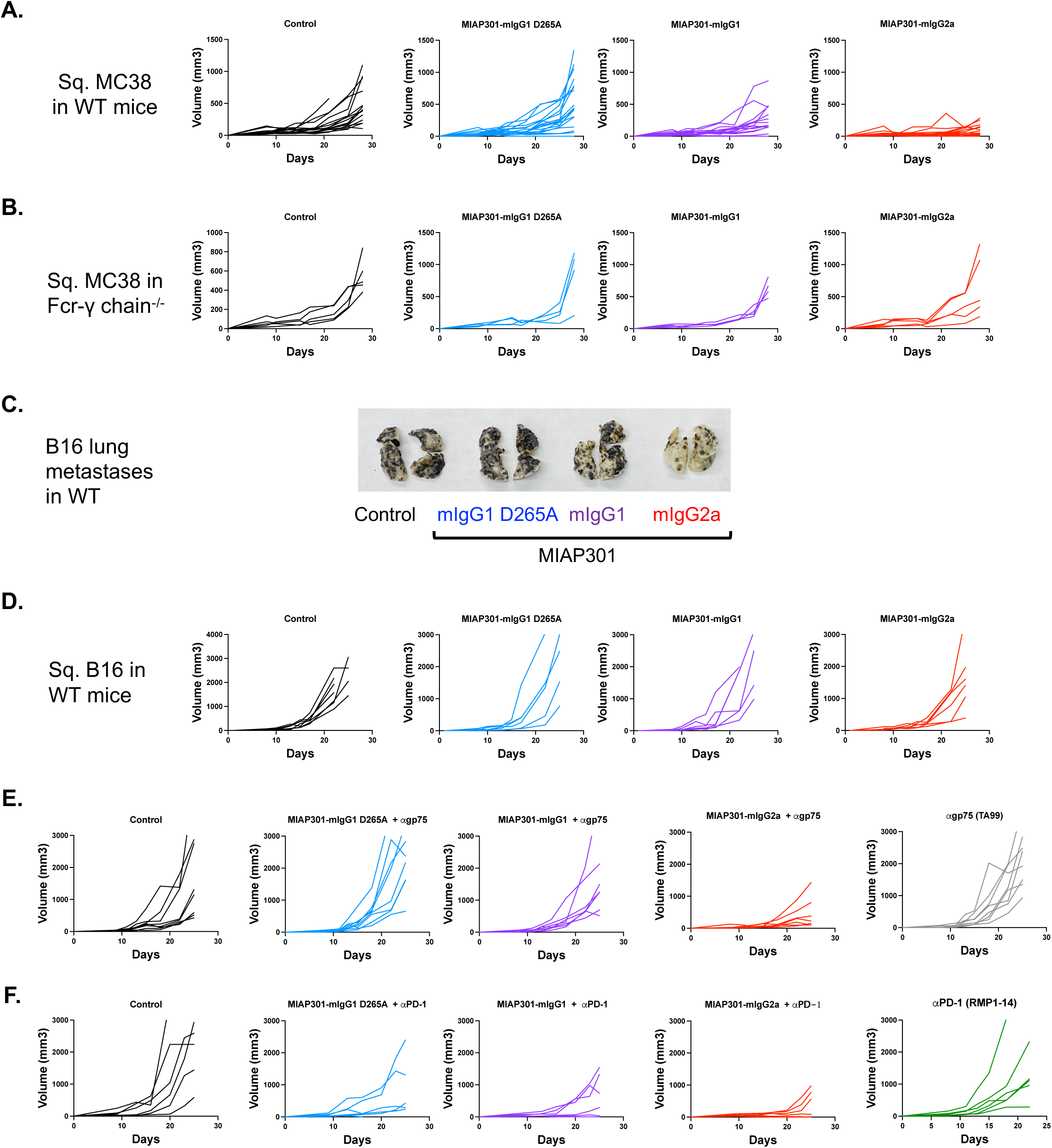
Engagement of activating FcγRs enhances in vivo antitumor activity of MIAP301 ab. **(A)** Individual growth of sq. MC38 tumors in WT mice, **(B)** in mice lacking all activating FcγRs (FcR-γ chain^-/-^) that were treated with MIAP301 Fc variants or control (50 ug IT, d. 8,10,14 and 18). **(C)** Representative images showing B16 lung metastases in WT mice after treatment with control or MIAP301 Fc variants (20 mg/Kg IP, d. 1,4,7 and 11). **(D)** Individual growth of sq. B16 tumors in WT mice, treated as monotherapy with MIAP301 Fc variants or control (50ug IT, d. 8,10,14 and 18); or **(E)** In combination with anti-gp75 ab (TA99-mIgG2a, 200 ug IP d. 8, 10, 14 and 18); or **(F)** In combination with anti-PD-1 ab (RMP1-14-mIgG1-D265A, 200 ug IP d. 8, 10, 14 and 18).

**Supplementary Figure 3.**
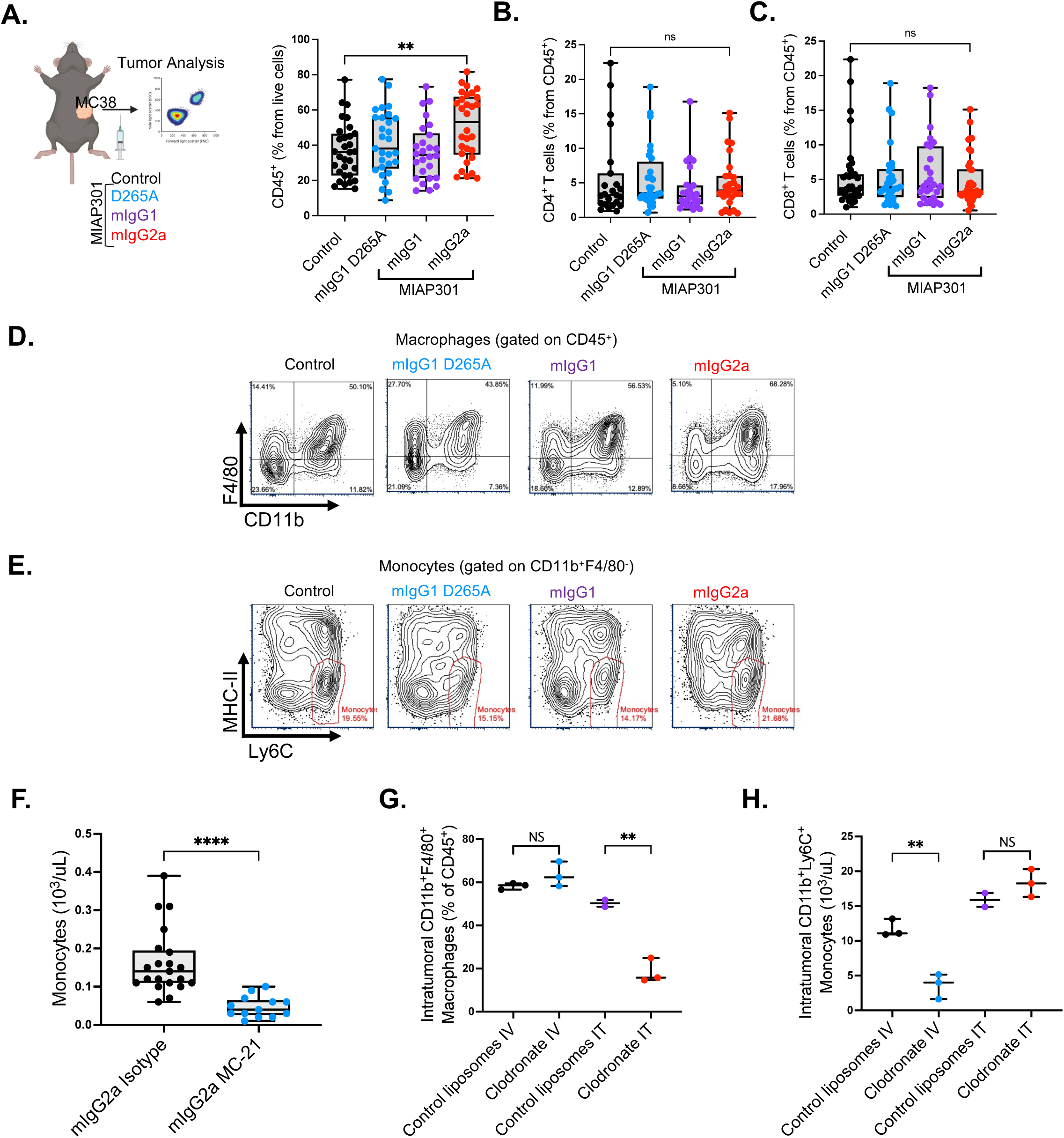
Changes myeloid and T cells after treatment with Fc variants of anti-mCD47 ab and assessment of depletion of monocytes and macrophages. **(A-C)**Established MC38 tumors were analyzed by flow cytometry 72 hours after treatment with Fc variants for CD47 antibodies (MIAP301). Quantification of CD45^+^, CD4^+^ and CD8^+^cells from MC38 tumors 72h after treatment with Fc variants is shown. **(D)** Representative flow cytometry plots of macrophages, and **(E)** monocytes 72h after treatment with MIAP301 Fc variants. **(F)** Peripheral blood levels of monocytes 24 hours after treatment with anti-CCR2 ab (Clone MC-21-mIgG2a, 100 ug) or isotype control in WT mice. **(G)** Quantification of intratumoral macrophages from MC38 tumors pre-treated with control- or clodronate-liposomes intravenously (IV) or intratumorally (IT) (100 uL, d.-3, -1) by flow cytometry. **(H)** Quantification of intratumoral monocytes from MC38 tumors pre-treated with control- or clodronate-liposomes intravenously (IV) or intratumorally (IT) (100 uL, d.-3, -1) by flow cytometry.

**Supplementary Figure 4.**
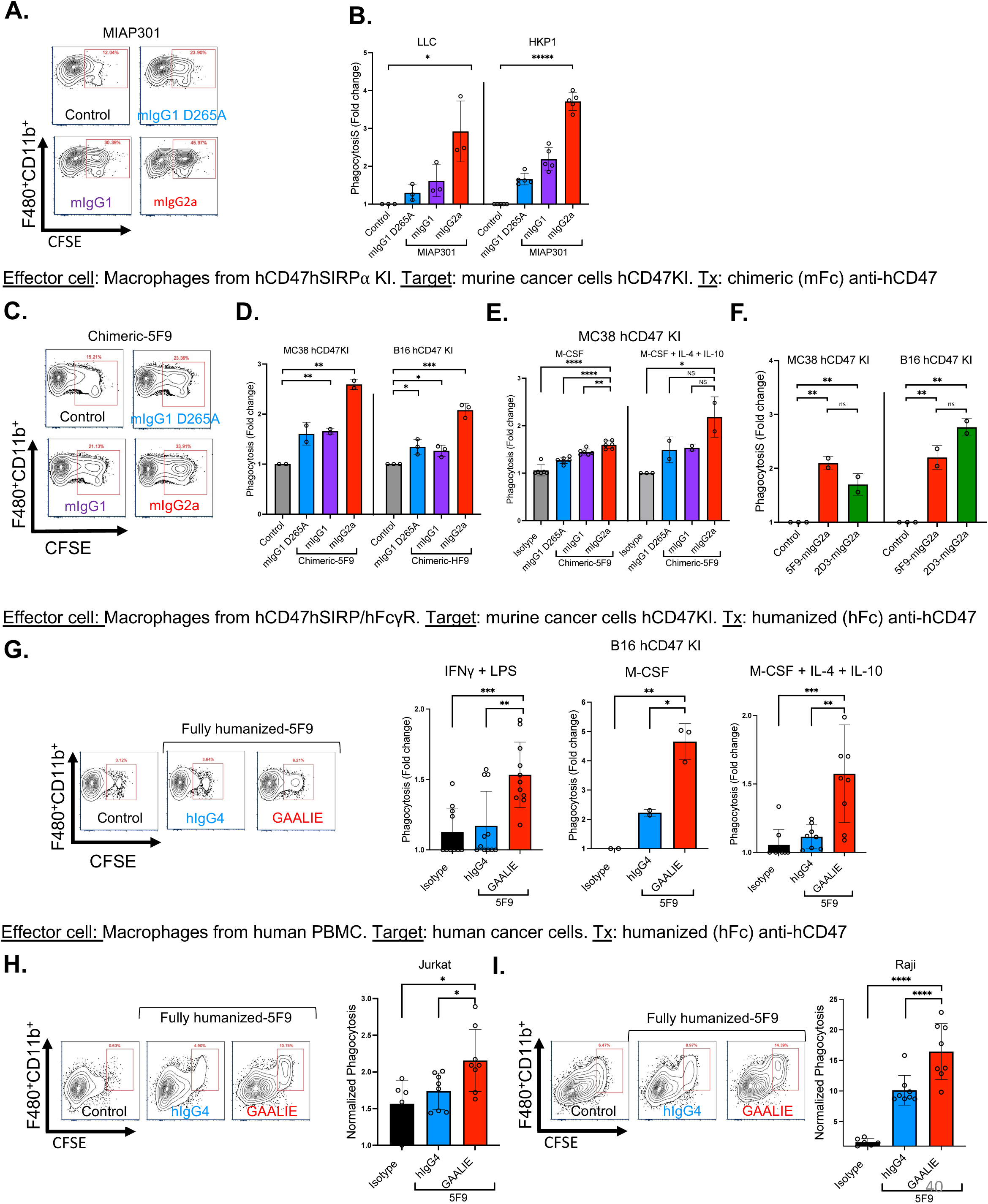
Fc-enhanced anti-CD47 antibodies enhance in vitro antibody dependent cellular phagocytosis (ADCP) across different species-match systems. **(A)** Representative flow cytometry plots of the percentage of phagocytosis of B16 cancer cells by BMDM from WT mice after treatment with control or Fc subclasses of MIAP301. **(B)** Phagocytosis of Lewis lung carcinoma (LLC) or HKP1 (Kras^G12D^p53^-/–^) lung adenocarcinoma after treatment with control or Fc subclasses of MIAP301. (**C)** Representative flow cytometry plot and **(D)** phagocytosis of MC38 hCD47 KI or B16 hCD47 KI by BMDM from hCD47/hSIRP⍺ KI mice after treatment with control or Fc variants of Chimeric-5F9. **(E)** Phagocytosis of MC38 hCD47 KI cells by BMDM from hCD47/hSIRP⍺ KI mice pre-treated with different polarization conditions. **(F)** Phagocytosis of MC38 hCD47 KI or B16 hCD47 KI cells by BMDM from hCD47/hSIRP⍺ KI mice after treatment isotype control, blocking (5F9), non-blocking (2D3) antibodies targeting hCD47, both in mIgG2a Fc format **(G)** (Left) Representative flow cytometry plots and (Right) phagocytosis of B16 hCD47 KI mice by BMDM from hCD47/hSIRP⍺/hFcγR mice after treatment with Fc variants of fully humanized anti-CD47 ab (5F9). Macrophages were pre-treated with different polarization conditions as indicated. **(H)** (Left) representative flow cytometry plots and (right) phagocytosis of human cancer cell line Jurkat and **(I)** Raji by human macrophages derived from PBMCs after treatment with Fc variants of fully humanized 5F9.

**Supplementary Figure 5.**
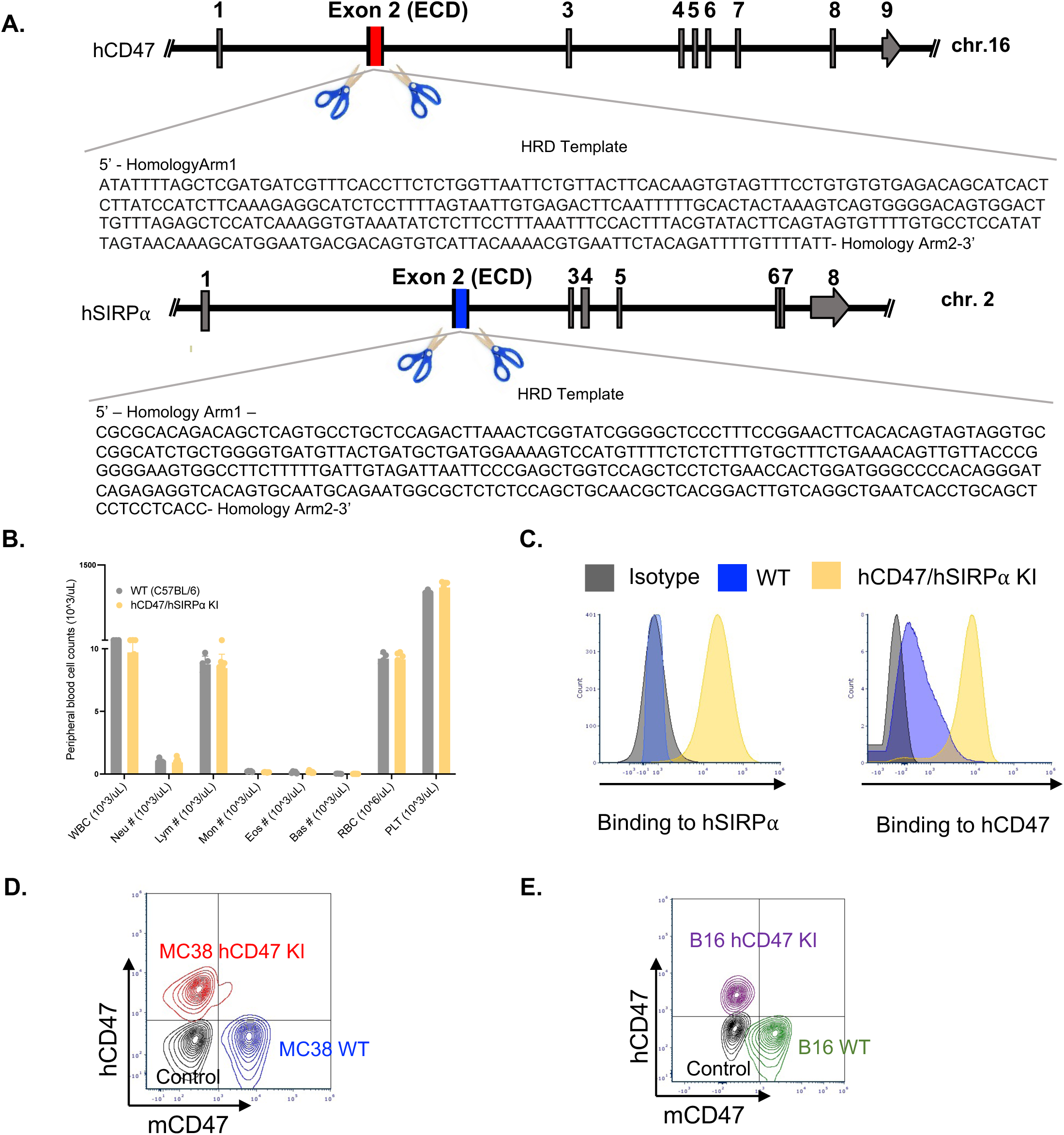
Development of humanized mouse model for CD47 and SIRPα (hCD47/hSIRPα KI mice) and cancer cell lines. **(A)** Schematic drawing showing CRISPR/Cas9-mediated gene-targeting strategy to generate CD47 (top) and SIRP⍺ (bottom) KI mice. The double-strand break induced by the CRISPR/Cas9 in the flanking regions of the ECD of mCD47 and mSIRP⍺ was replaced by their human counterparts via non-homologous end joining. **(B)** Complete cell blood count of WT and hCD47/hSIRP⍺ KI mice **(C)** Right: Recombinant hSIRPα was incubated with RBCs from WT and hCD47/hSIRPα KI mice, binding was only detected in RBCs from the hCD47/hSIRPα mouse. Left: Recombinant hCD47 was incubated with leukocytes from WT and hCD47/hSIRPα KI mice, binding was primarily detected in leukocytes from the hCD47/hSIRPα mouse. **(D)** Flow cytometry analysis of expression of hCD47 and mCD47 in unstained (Black), MC38 WT (blue) and MC38 hCD47 KI cells (red). **(E)** Flow cytometry analysis of expression of hCD47 and mCD47 in unstained (Black), B16 WT (blue) and B16 hCD47 KI cells (red).

**Supplementary Figure 6.**
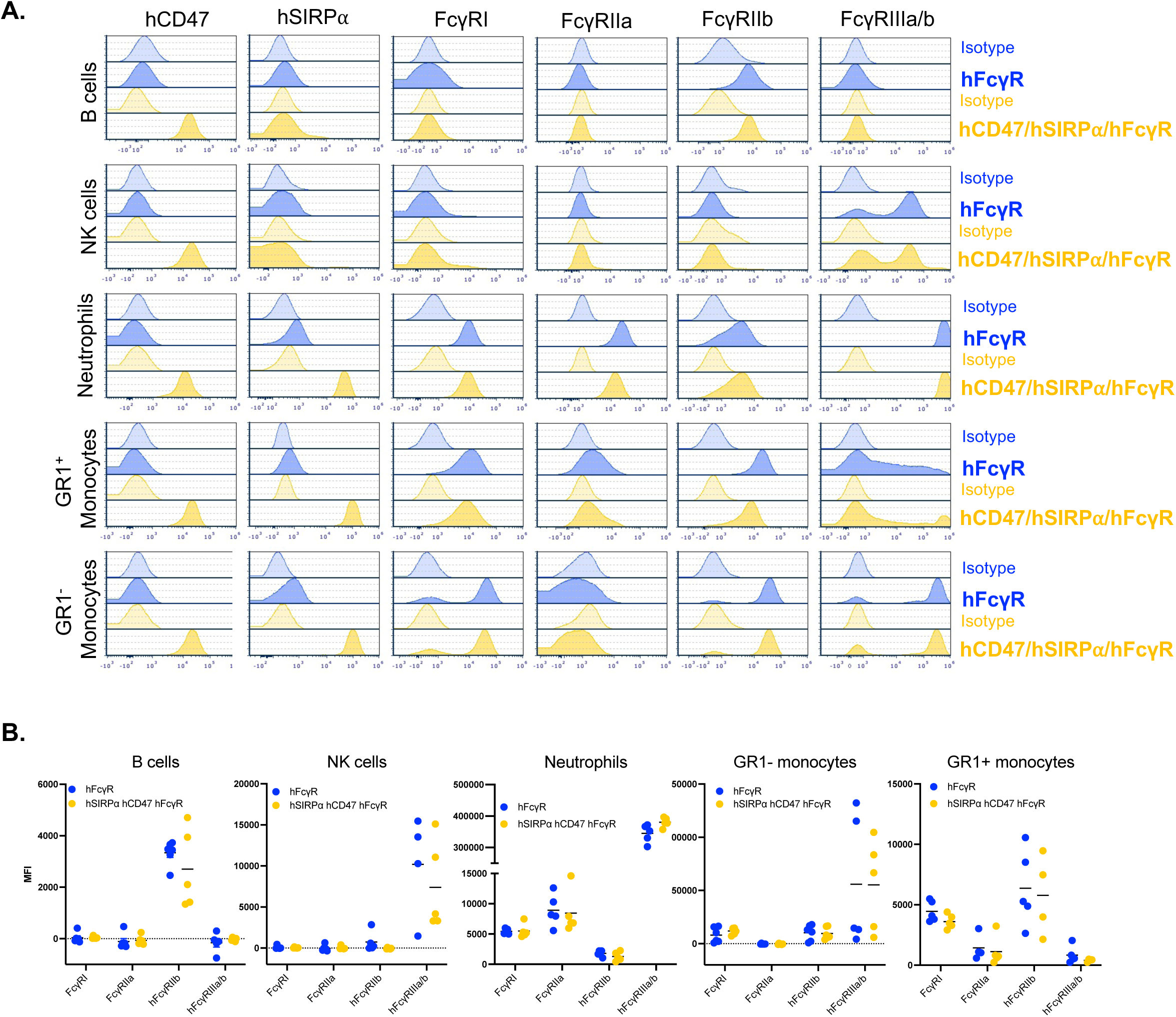
Characterization of FcγRs in mice humanized for CD47, SIRP⍺ and FcγRs. **(A)** Characterization of FcγR profile on mice humanized for FcγRs (blue) and mice humanized for CD47, SIRP⍺ and FcγRs (yellow), representative flow cytometry plots and **(B) Q**uantification are shown.

